# BRD4-mediated epigenetic regulation of endoplasmic reticulum-mitochondria contact sites is governed by the mitochondrial complex III

**DOI:** 10.1101/2024.02.02.578646

**Authors:** Brandon Chen, Drew C. Stark, Pankaj V. Jadhav, Theophilus M. Lynn-Nguyen, Benjamin S. Halligan, Nicholas J. Rossiter, Nicole Sindoni, Myunsun Shin, Joao A. Paulo, Matthew Chang, Imhoi Koo, Sergei Koshkin, Sanjana Eyunni, Paolo Ronchi, Michelle T. Paulsen, Pietro Morlacchi, David A. Hanna, Jason Lin, Rachel M. Guerra, David J. Pagliarini, Ruma Banerjee, Abhijit Parolia, Mats E. Ljungman, Andrew D. Patterson, Joseph D. Mancias, Shyamal Mosalaganti, Jonathan Z. Sexton, Tito Calì, Costas A. Lyssiotis, Yatrik M. Shah

## Abstract

Inter-organellar communication is critical for cellular metabolic homeostasis. One of the most abundant inter-organellar interactions are those at the endoplasmic reticulum and mitochondria contact sites (ERMCS). However, a detailed understanding of the mechanisms governing ERMCS regulation and their roles in cellular metabolism are limited by a lack of tools that permit temporal induction and reversal. Through unbiased screening approaches, we identified fedratinib, an FDA-approved drug, that dramatically increases ERMCS abundance by inhibiting the epigenetic modifier BRD4. Fedratinib rapidly and reversibly modulates mitochondrial and ER morphology and alters metabolic homeostasis. Moreover, ERMCS modulation depends on mitochondria electron transport chain complex III function. Comparison of fedratinib activity to other reported inducers of ERMCS revealed common mechanisms of induction and function, providing clarity and union to a growing body of experimental observations. In total, our results uncovered a novel epigenetic signaling pathway and an endogenous metabolic regulator that connects ERMCS and cellular metabolism.

## Main Texts

The dynamic regulation of organelle networks and inter-organellar communication is critical for cellular metabolism and physiology. Direct inter-organellar communication occurs at membrane contact sites, which directly regulate metabolite transport, protein complex organization, signaling and organellar function^1,2^. One of the most well studied and abundant inter-organellar contact sites are <u>e</u>ndoplasmic reticulum-<u>m</u>itochondria <u>c</u>ontact <u>s</u>ites (ERMCS). While a wealth of research has defined the unique proteome at ERMCS, the precise spatiotemporal regulation by various signaling pathways, and evolutionarily conserved molecular tethering complexes that structurally support ER-to-mitochondria interactions remains still elusive. Functional genetic and biochemical studies have revealed essential roles of ERMCS, including phospholipid synthesis^3^, calcium buffering^4^, mitochondrial dynamics^5,6^, mitochondrial DNA distribution^7^ and autophagosome formation^8^.

Dysregulation of ERMCS and tethers contributes to the etiology of various disease, including neurodegeneration^9^, obesity^10^, cancer^11^, diabetes^12^, and inborn errors of metabolism^13^. Understanding the fundamental mechanisms underlying ERMCS dysregulation is essential for identifying potential drug targets to restore ERMCS function in disease. However, significant gaps persist in our basic understanding of the molecular drivers governing the remodeling and organization of ERMCS. Moreover, the exact mechanisms by which the microenvironment, signal transduction, and molecular cues initiate, maintain, and modify adaptive responses of ERMCS are yet to be fully elucidated.

In this study, we have identified a novel epigenetic response and a metabolic redox state that regulate the formation and dynamics of ERMCS. We characterized an FDA-approved compound, fedratinib, which induces ERMCS formation via BRD4-mediated histone recognition and described it as a new tool for temporal, reversible control of ERMCS. Fedratinib treatment establishes selective ER wrapping around mitochondria with cristae structure defects, membrane potential loss, and metabolic rewiring. ERMCS induction requires an intact mitochondrial complex III, which is a major site of coenzyme Q (CoQ) oxidation. Attempts to modulate mitochondrial CoQ redox state suggest a potential mechanism where an increase in the reduced-to-oxidized CoQ ratio could block ERMCS. Importantly, our study demonstrates that the transcriptional and metabolic requirements for ERMCS induction by fedratinib are common to other cellular stressors known to triggers ERMCS and underscores the existence of a conserved regulatory network governing ERMCS.

### High throughput pharmacogenomic screening identifies novel regulators of ERMCS

To follow ERMCS in living cells in real-time, we generated a panel of eleven isogenic gene-edited cell lines with a novel reversible split fluorescent ERMCS reporter, termed <u>spl</u>it-GFP based <u>c</u>ontact site <u>s</u>ensor (SPLICS) ^14–16^. This genetically encoded sensor is stably integrated with a doxycycline-inducible promoter for titrating reporter expression^17^. In addition, we engineered a mitochondria matrix reporter (mitoTagRFP) to control for gene expression and monitor mitochondrial abundance^18^ (Fig. 1a). Single cell clones were profiled for those with inducible and reversible green fluorescence as well as stable red fluorescence. Probe induction did not have a deleterious effect on cell growth for 7 days, nor on oxygen consumption (OCR) or extracellular acidification rate (ECAR) at 72 hours post reporter induction (Extended Data Fig. 1a and b). To demonstrate the sensitivity, accuracy, and specificity of the probe, SPLICS lines were treated with a known inducer of ERMCS, the ER stress inducer thapsigargin^19^ (Fig 1b), or by over-expressing an artificial ER-mitochondria linker (ER-MT tether). Both treatments increased the SPLICS signal significantly (Fig. 1c). Moreover, using control pairs of splitGFP probes: mKate2-β_11_ + cytosolic GFP_1-10_, mKate2-β_11_ + outer mitochondria membrane (OMM) GFP_1-10,_ ER_short_ + cytosolic GFP_1-10_, we did not see a remarkable increase in any of these probe pairs in cells treated with thapsigargin^20^ (Extended Data Fig. 1e). Using this inducible SPLICS system, we generated eleven isogenic human and mouse cell lines in investigate ERMCS regulation.

**Fig. 1:**
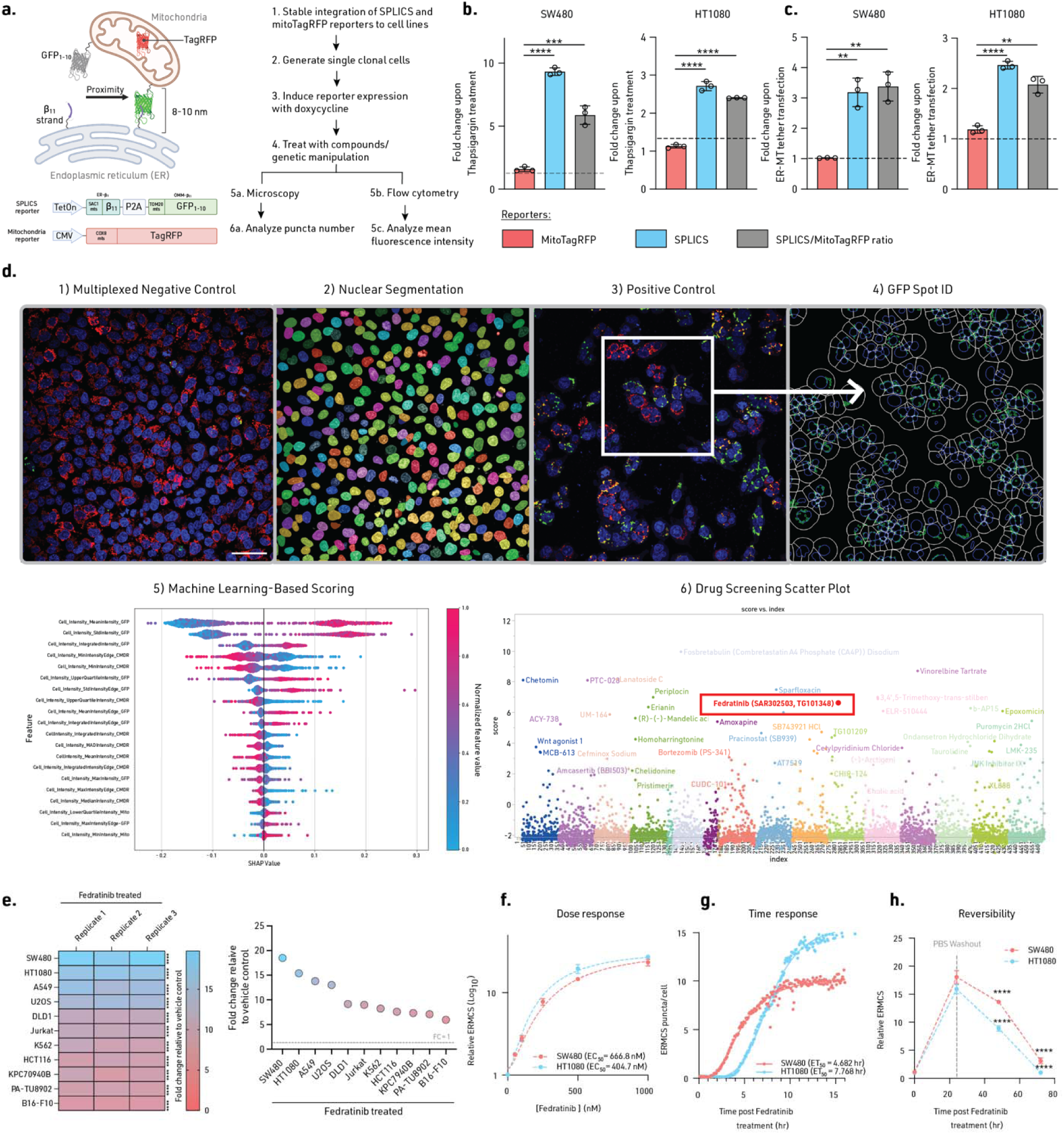
High throughput image-based chemical screen identifies novel regulators of ERMCS. **a**, Schematic of the SPLICS and mitochondria TagRFP (mitoTagRFP) reporter and analysis pipeline for ERMCS. **b**, Relative fluorescence intensity of SPLICS, mitoTagRFP, and SPLICS/mitoTagRFP ratio following treatment with vehicle or thapsigargin (50 nM) for 24 hr, or **c**, 48 hr post transfection of ER-MT tether. **d**, D. Schematic representation of the bioassay design for the ERMCS drug screen. Blue: Hoechst 33342/nuclei. Red: MitotagRFP/mitochondria. Green: SPLICS/ERMCS. Clockwise: 1. Negative control demonstrating low levels of ERMCS; 2. Cellpose nuclear segmentation using Hoechst counterstain; 3. Positive control demonstrating induction of high levels of ERMCS. 4. GFP spot identification and feature extraction using Cellprofiler for ERMCS. Machine learning was performed by training an XGBoost model against per-plate controls. 5. Shapley additive explanations showing GFP features used to identify hits in drug screening. Beeswarm plot showing SHAP values indicating feature importance for drug effect scoring based on GFP features. 6. Drug screening scatter plot showing drug tested on the x-axis and XGBoost score on the y-axis indicating ERMCS induction. Each color categories for individual data points suggest different plates instead of categorization of drug pathways. Fedratinib is highlighted in red on the scatter plot. Scale bar: 15 μm. **e**, Relative ERMCS in a panel of cell lines treated with vehicle or fedratinib treatment for 24 hr. **f**, Relative ERMCS of cells treated with vehicle or fedratinib for 24 hr, followed by PBS washout, and monitored for additional 48 hr. **g**, Dose dependent increase of relative ERMCS of cells treated with vehicle or fedratinib for 24 hr. **h**, Time-lapse imaging using Lattice light sheet microscopy to monitor ERMCS puncta of cells treated with fedratinib for 16 hr with quantified puncta per cell. Experiment (**b, c, e, f, h**) were performed three times, and experiment (**d**) was performed once, and experiment (**g**) was performed twice. Significance was calculated with an unpaired two-tailed t-test. *P ≤ 0.05, **P ≤ 0.01, ***P ≤ 0.005, ****P ≤ 0.001. Data are mean ± SD. Statistical source data are provided in Source Data Fig. 1.

To discover novel mediators and chemical tools of ERMCS induction, we developed a high-content phenotypic imaging screen with machine learning categorization approaches using our SPLICS lines and assayed an FDA-approved drug repurposing library (Fig. 1d). Consistent with known roles of ER stress in ERMCS, we identified proteasome inhibitors bortezomib and carfilzomib^21^. Moreover, we identified microtubule inhibitors that modulate organelle movement within in cells^22^ (Supplementary Table 1 and Zenodo DOI: <u>10.5281/zenodo.13963526</u>). We decided to focus on fedratinib as it induced high levels of ERMCS, and its annotated mechanisms of action have not been associated with ERMCS regulation (Fig. 1d). Fedratinib inhibits Janus kinase 2 (JAK2) as well as bromodomain-containing protein 4 (BRD4)^23^. The increase in ERMCS following fedratinib treatment was observed in a large panel of human and mouse cell lines (Fig 1e). We focused on SW480 and HT1080, as these had the highest fold change by fedratinib. ERMCS induced by fedratinib was both dose- and time-dependent with estimated EC_50_ and ET_50_ values of 666.8 nM/4.7hr (SW480) and 404.7 nM/7.8 hr (HT1080, which was evident using both the SPLICS reporter lines as well as by an orthogonal proximity labeling method (Fig. 1f and g; Extended Data Fig. 1h, Supplementary Movies 1 and 2). Importantly, the increase in ERMCS was reversible following a washout of fedratinib from the culture media, highlighting the capability of SPLICS reporters to dynamically and reversibly measure organelle contact sites in living cells (Fig. 1h). In addition, fedratinib did not significantly increase fluorescence of our control splitGFP probe pairs, nor did it induce autofluorescence when SPLICS is not activated (Extended Data Fig. 1f). Collectively, this screen identified the first FDA-approved drug with potent bioactivity in increasing ERMCS across cell lines with distinct tissue and organisms of origin.

### Bromodomain and Extra-Terminal protein (BET)-dependent transcription response is required for ERMCS induction

Fedratinib was first described as a JAK2-specific inhibitor for treating myelodysplastic syndromes^24^. Thus, we tested the ERMCS-inducing activity of other JAK2 inhibitors. We found that the JAK2 inhibitor Ruxolitinib did not induce ERMCS formation at a concentration that inhibits JAK2-dependent STAT3 phosphorylation (Fig. 2a and Extended Data Fig. 2a). In addition, cells with CRISPR interference (CRISPRi) mediated JAK2 knockdown (KD) still responded to fedratinib-dependent ERMCS induction (Extended Data Fig. 2c and d). Therefore, both genetic and pharmacological approaches suggest JAK2 is not the main target of fedratinib for inducing ERMCS.

**Fig. 2:**
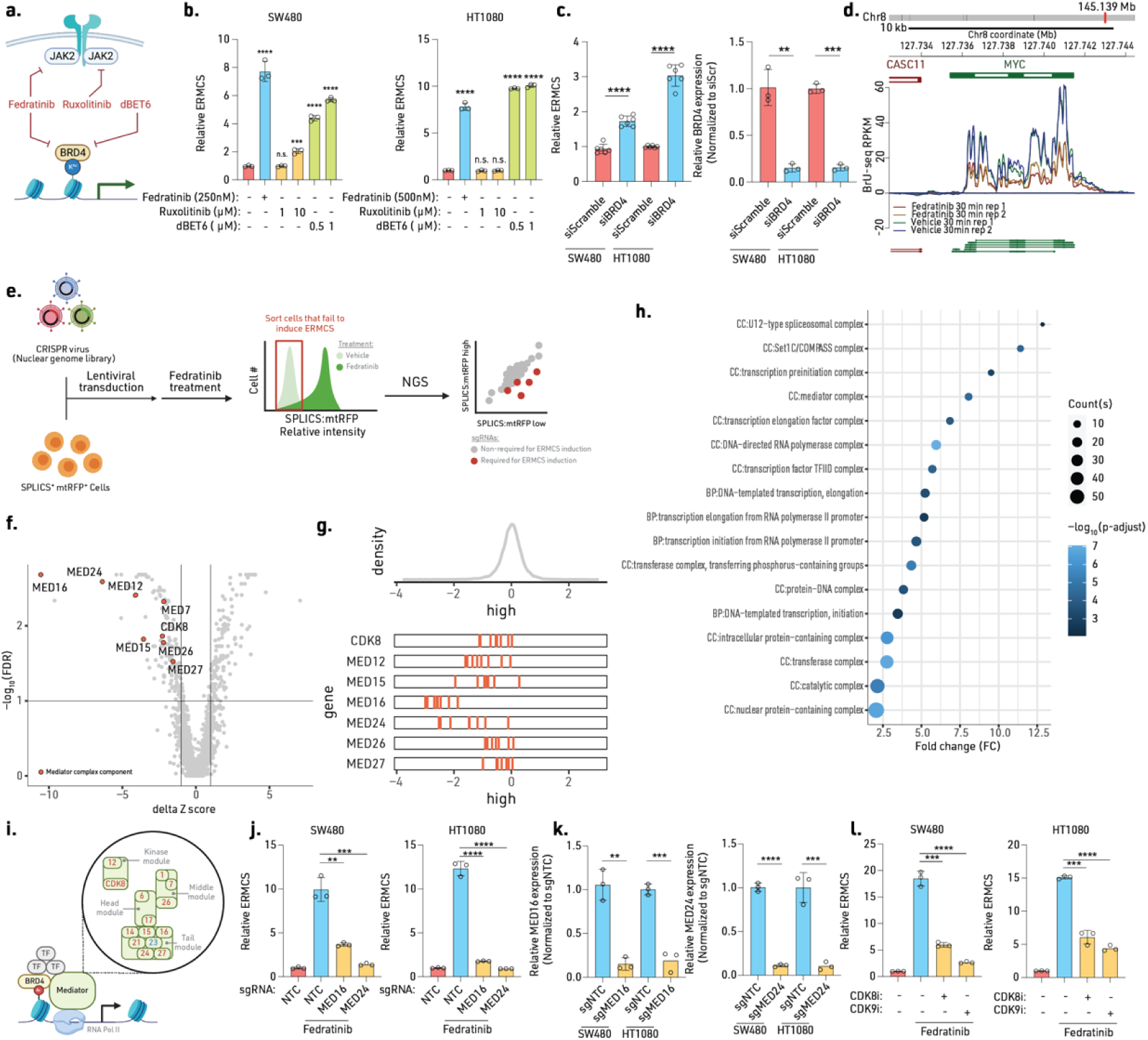
ERMCS induction requires a transcriptional response dependent on bromodomain and extra-terminal protein (BET). **a**, Mechanisms of action for fedratinib, ruxolitinib, and dBET6 targeting JAK2 and BRD4 (Left). Relative ERMCS in cells treated with JAK2 and/or BRD4 inhibitors. **b**, Relative ERMCS and BRD4 mRNA expression of cells following knockdown (KD) using scrambled or BRD4 targeting siRNAs. **c**, Relative ERMCS in cells treated with inhibitors of transcription kinase CDK8 (AS-2863619, 1 μM) or CDK9 (NVP2, 1 μM) for 6 hr, and then treated with vehicle or fedratinib for 24 hr. **d**, Representative BrU-seq transcript profile of SW480 cells treated with fedratinib or vehicle for 30 min. **e**, Schematic of a fluorescence-activated cell sorting (FACS)-based CRISPR screen using a nuclear genome targeted sgRNA library. **f**, Volcano plot of BrU-seq analysis after SW480 cells treated with fedratinib for 2 hr (top panel). X-axis is the CRISPR gene scores (delta Z score) with Y-axis being -Log_10_ of false discovery rate (FDR). Highlighted red circles are sgRNAs targeting the Mediator complex components. **g**, Histogram indicating the distribution of sgRNA. Shown below is the relative enrichment of mediator complex genes. **h**, GSEA ridgeplot displaying significantly enriched (FDR < 0.01) protein complexes based on differential CRISPR screen scores.) BP: Biological processes. CC: Cellular components. **i**, Schematic of the mediator complex and BRD4 in transcriptional regulation. Mediator components from the CRISPR library, annotated in red demonstrate suppression of ERMCS. **j**, Relative ERMCS and gene expression level of MED16 and MED24 mRNA transcripts upon CRISPRi mediated knockdown of cells treated with vehicle or fedratinib for 24 hr. **k**, Relative MED16 and MED24 expression of sgMED16 and sgMED24 cells. Experiments. **l**, Relative ERMCS in cells treated with inhibitors of transcription kinase CDK8 (AS-2863619, 1 μM) or CDK9 (NVP2, 1 μM) for 6 hr, and then treated with vehicle or thapsigargin (50 nM) for 24 hr. (**b**, **c**, **j**-**l**) were performed three times, experiment (**d**) was performed once with duplicates, and experiment (**e**-**h)** was conducted once. Significance was calculated with an unpaired two-tailed t-test. n.s. = not significant, *P ≤ 0.05, **P ≤ 0.01, ***P ≤ 0.005, ****P ≤ 0.001. Data are mean ± SD. Statistical source data are provided in Source Data Fig. 2.

Fedratinib has other targets, including BRD4, a member of the bromodomain and extra-terminal (BET) epigenetic reader family^25–27^. Fedratinib binds to the acetylated-lysine recognition pocket of BRD4 and prevents BRD4 interaction with acetylated histones. Indeed, we observed that canonical BRD4-hypersensitive targets, such as c-MYC, were decreased in expression upon fedratinib treatment (Extended Data Fig. 2b and m). Similar to fedratinib, other inhibitors of BRD4 from our small molecule screen also increased ERMCS (i.e. OTX-015 and IBET-762; Zenodo DOI: <u>10.5281/zenodo.13963526</u>.). To directly assess BRD4, we utilized the PROTAC degrader of BRD4, dBET6, and siRNA-mediated KD, both of which increased ERMCS (Fig. 2a, b, and c; Extended Data Fig. 2a).

Next, we directly investigated whether both fedratinib and thapsigargin works through BRD4 inhibition. First, we conducted chromatin fractionation assay and found that fedratinib and the positive control BET inhibitor JQ1 displaced BRD4 from chromatin, but not thapsigargin (Extended Data Fig. 2d). This suggests that fedratinib inhibits BRD4 function by directly altering its chromatin occupancy, but thapsigargin does not. Second, we investigated how fedratinib and thapsigargin differentially affect genome wide BRD4 occupancy via chromatin immunoprecipitation (ChIP). BRD4 ChIP-seq conducted from SW480 cells identified that fedratinib has fewer BRD4 bound genomic loci in both promoter and non-promoter regions than thapsigargin (Extended Data Fig. 2f and g), and the BRD4 bound regions are mostly promoters, introns, or intergenic regions (Extended Data Fig. 2h). We then confirmed with ChIP-qPCR in HT1080 cells that fedratinib led to loss of BRD4 occupancy at the PVT enhancer regions that controls BRD4-hypersensitive gene MYC (Extended Data Fig. 2i). Therefore, we confirm fedratinib acts through BRD4 inhibition to induce ERMCS while thapsigargin’s mechanism is BRD4 independent.

To identify whether BRD4-dependent ERMCS induction is transcriptionally regulated, we inhibited transcription kinases cyclin-dependent kinases 8 and 9 (CDK8 and CDK9). Inhibition of CDK8 and CDK9 both suppressed ERMCS induction (Fig. 2l and Extended Data Fig. 2e), thus pointing out the importance of an active transcriptional program for ERMCS regulation. To further test this we performed a pulse-chase experiment with RNA polymerase II (RNA pol II) inhibitor Actinomycin D (ActD). Fedratinib- and thapsigargin-induced ERMCS were suppressed when transcription activity was rapidly blocked (Extended Data Fig. 2f and g), whereas the mitochondrial RFP signal was stable following acute transcriptional blockade.

To characterize the early transcriptional response upon ERMCS induction, we conducted bromouridine (BrU) labeling of nascent mRNA sequencing, BrU-Seq, at 30 min and 2 hr after fedratinib treatment (Supplemental table 3). BRD4 hypersensitive target gene MYC decreased its nascent mRNA transcription within 30 mins of fedratinib treatment (Fig. 2d). Through gene set enrichment analyses (GSEA), we identified potent upregulation of genes associated with the integrated stress response (ISR) and the unfolded protein response (UPR). In addition, many of the ISR genes were increased at the protein level when measured by quantitative proteomics (Extended Data Fig. 3a, b, and c and Supplementary Table 2). While thapsigargin is thought to increase ERMCS via inducing ER stress, fedratinib did not increase UPR markers phospho-PERK, IRE1α, CCPG1, ATF6, spliced XBP1, CHOP and BiP (Extended Data Fig. 5a). Moreover, inhibiting ER stress propagation with chemical chaperone tauroursodeoxycholic acid (TUDCA) did not reverse ERMCS induction (Extended Data Fig. 5d). Therefore, we believe the induction of ERMCS by fedratinib is independent of the UPR.

However, we observed increased levels of ISR markers ATF4 and phosphorylated-eIF2α at Ser51 upon fedratinib treatment (Extended Data Fig. 3d). To identify which eIF2α kinase arm drives this ISR signature, we treated our cells with inducers of eIF2AK1/2/3/4 (GCN2, PKR, PERK, HRI). The most robust inducer of ERMCS was the PKR activator BEPP, and inhibition of PKR with its inhibitor PKR-IN-C16 reversed ERMCS induction and ATF4 expression (Extended Data Fig.3e, g, h, and i). ISR inhibitor ISRIB decreased ATF4 expression by thapsigargin and fedratinib treatment, yet it did not suppress ERMCS (Extended Data Fig. 3f). We, therefore, posit that fedratinib induces ERMCS via PKR activation, but the ISR is unlikely to be a direct activator of ERMCS. Additionally, fedratinib did not activate UPR^mito^, and activating mitochondrial ISR via oligomycin or CCCP was not sufficient to induce ERMCS formation (Extended Data Fig. 3k and 6c).

We also observed that genes associated with autophagy increased with fedratinib treatment. BRD4 inhibition has been shown to induce autophagy^28^, and autophagosomes have been shown to emerge from ERMCS^8^. Indeed, we confirmed increased autophagy with fedratinib treatment by immunoblotting for lipidated LC3B (Extended Data Fig. 4a). Additionally, when we combined fedratinib with an autophagy inhibitor chloroquine (CQ), we observed enhanced LC3B lipidation, suggesting an elevated autophagic flux (Extended Data Fig. 4a). Fedratinib treatment also increased lysosomes measured by lysotracker intensity in a dose dependent manner (Extended Data Fig. 4b). However, even after inhibition of autophagy with CQ or V-ATPase pump inhibitor Bafilomycin A (BafA), we still observed ERMCS formation (Extended Data Fig. 4c). Moreover, colocalization studies of lysosomes and autophagosomes with SPLICS did not show overlap at ERMCS (Extended Data Fig. 4d and e), indicating ERMCS was not an early step of mitophagosome formation. Therefore, these data suggest autophagy induction is a result of fedratinib treatment but not an inducer of ERMCS formation.

To identify the BRD4 co-regulated complex and transcription factors that coordinate ERMCS modulation, we performed an unbiased CRISPR screen coupled with fluorescent activated cell sorting (FACS), using a nuclear genome library. Following transduction with the CRISPR library, which contained genes annotated with nuclear localization or roles in epigenetic modification, we induced ERMCS formation with fedratinib, and sorted cells that failed to induce ERMCS (bottom 10% SPLICS^low^/mitoTagRFP) (Fig. 2e). Through gene set enrichment analyses (GSEA), we found a potent signature from components of the mediator complex (Fig. 2f, g, and h; Supplementary Table 4). The mediator complex is a hetero-multimeric protein complex that recruits other transcription factors, co-regulators, and RNA pol II machineries along with BRD4. We generated knockdowns of mediator complex components MED16 and MED24 KD via CRISPR interference (CRISPRi), which failed to induce ERMCS in response to fedratinib or thapsigargin (Fig. 2j and k and Extended Data Fig. 2j and k).

Taken together, here we have demonstrated that Fedratinib induces ERMCS via suppression of BRD4, and this activity is transcriptionally controlled. Moreover, induction of ERMCS via the ER stressor thapsigargin also depends on an active gene expression program, indicating that transcriptional rewiring to be common for other modes of ERMCS induction.

### Fedratinib induces ER-wrapping of mitochondria at ERMCS

Organelle structures and inter-organellar organization networks inform the metabolic state of cells. An increase in ERMCS drives adaptive oxidative phosphorylation and supports ER homeostasis. Therefore, we postulate that understanding how fedratinib impacts ERMCS architecture will provide valuable information for a deeper understanding of the metabolic factors, regulation, and consequences of ERMCS induction. To address this, we utilized various electron microscopy imaging approaches to sequentially characterize the ERMCS architecture (Fig 3a).

**Fig. 3:**
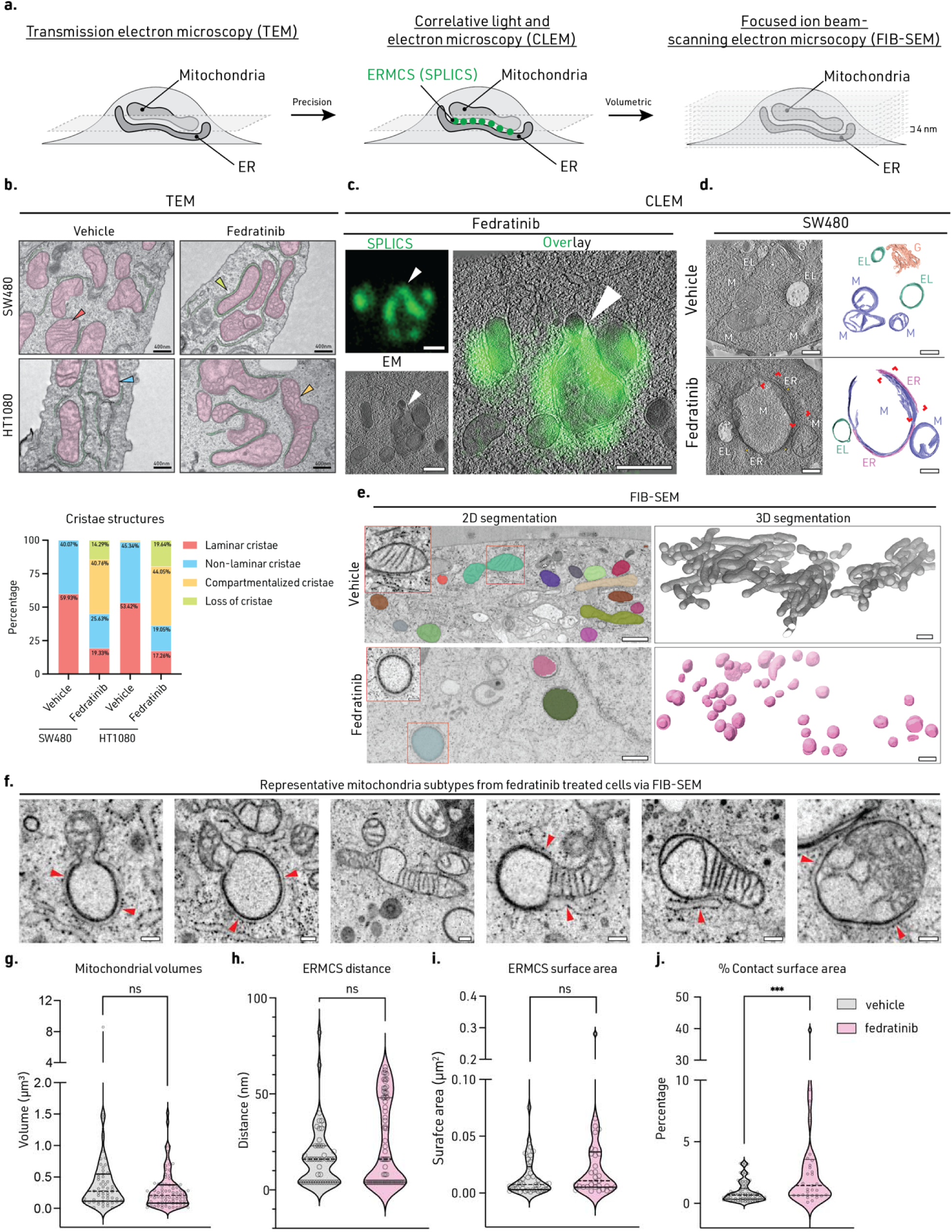
Fedratinib induces ER-wrapping of mitochondria at ERMCS. **a**, Workflow of our multi-modal electron microscopy imaging approach. **b**, Representative TEM images (scale bar: 400 nm) of WT cells treated with vehicle or fedratinib for 24 hr (Mitochondria green, ER: magenta (Above)). Distribution of cristae morphologies quantified from 25 cells as a percentage of total mitochondria (n=161-299) (Below). Cristae structures are annotated in corresponding colors in the stacked bars with percentage of each classification listed. Red: Laminar cristae. Blue: Non-laminar cristae. Yellow: Compartmentalized cristae. Green: Loss of cristae. Each example cristae structure is indicated with its corresponding color in the EM images. **c**, Correlative light and electron microscopy (CLEM) images of SW480 cells treated with fedratinib for 24 hr (Scale bar: 500 nm). White arrows indicate an example of SPLICS^Hi^ area. **d**, Segmentation of organelle membranes of SW480 cells treated with fedratinib for 24 hr (M: mitochondria, ER: endoplasmic reticulum, EL: endolysosome, G: golgi apparatus; Scale bar: 500 nm). **e**, Representative slices through the tomograms of vehicle control (top left) and fedratinib-treated (bottom left) HT1080 cells with an overlay of 3D segmentation. Inset highlights a representative mitochondrion in each case. Scale bar, 200 nm and 50 nm for inset. 3D volume reconstructions(right) displaying mitochondria. Scale bars, 1 μm. **f**, FIB-SEM images showing instances of different morphologies of mitochondria wrapped around by ER membrane (red arrowheads). Scale bar, 200 nm. **g**, Quantification of total mitochondrial volume. **h**, ER-mitochondria contact site intermembrane distances. **i**, ER-mitochondria contact surface area. **j**, Percentage contact surface area for individual mitochondria in vehicle-treated and fed-treated cells. Experiments (**a-j)** were performed once. Red rectangle highlights area with cristae collapse. Statistical source data are provided in Source Data Fig. 3.

From our transmission electron microscopy (TEM) on SW480 and HT1080 cells. We observed structural changes, such as the ER membrane enveloping the mitochondria and the collapse of cristae, we also noted the compartmentalization of structures within the inner mitochondrial membrane (IMM) (Fig. 3b). In addition, approximately 15-20% of mitochondria exhibited complete loss cristae and 40-45% demonstrated compartmentalization of the IMM (Fig. 3b). Although we did not observe consistent changes in circular cross-section, area, or aspect ratio, a higher variability of these structural indices were seen in fedratinib-treated cells (Extended Data Fig. 8d).

To further corroborate our observations and specifically probe ERMCS, we performed correlative light and electron microscopy (CLEM) using the SPLICS reporter that precisely marks ERMCS. We collected tilt-series guided by fluorescence from ERMCS in SW480 cells reconstructed a three-dimensional tomogram (Fig. 3c and d). Fedratinib-treated cells, but not vehicle controls, exhibited direct juxtaposition of the ER and outer mitochondrial membrane (OMM) at specific SPLICS^Hi^ locations (Fig. 3d), confirming our initial TEM observations (Fig. 3b). Furthermore, we observed near-complete collapse of the inner mitochondrial membrane/cristae at these contact sites. Typically, ERMCS are characterized by segments of the ER contacting the OMM rather than complete ER envelopment of mitochondria. However, following fedratinib treatment, we observed striking ER-mitochondria engulfment. Thus, fedratinib treatment not only induced enhanced ER-mitochondria contact but also promoted unanticipated ultrastructural reorganization at these sites.

Our TEM and CLEM approaches revealed ER-enclosure structures around select mitochondria. To obtain global higher-resolution volumetric information of ERMCS, we used high pressure freezing succeeded by freeze-substitution and focused ion beam-scanning electron microscopy (FIB-SEM) on fedratinib- or vehicle-treated HT1080 cells. This approach preserves cellular components in their near native state, avoiding chemical fixation artefacts that might disrupt their delicate organization. Imaging through half a cell volume at 4 nm isotropic pixel size. We trained a deep-learning model to distinguish mitochondria and ER. We segmented these organelles in our dataset and identified mitochondria ensconced by ER in fedratinib-treated cells (Fig. 3e). Consistent with our light microscopy data, we observed a range of mitochondrial and cristae structure with varying degrees of ERMCS in fedratinib-treated cells (Fig. 3f). Overall, we did not observe significant global changes in mitochondrial volume, ERMCS distance, or ERMCS surface area (Fig. 3g-i). However, analysis of individual mitochondria revealed a substantial increase in percent contact area (Fig. 3j). In conclusion, by combining multiple EM modalities, we identified a novel ER-wrapped mitochondrial morphology in fedratinib-treated cells, characterized by a heterogeneous mitochondrial population with varying degrees of ERMCS.

### Metabolic and lipidomic changes associated with selective depolarization of ER-wrapped mitochondria

To visualize ERMCS in a larger cellular context, intact mitochondria and ER networks, dynamics, and morphology in the whole cell were imaged with light microscopy (LM). We found that a subset of mitochondria were rounded and swollen in fedratinib-treated cells, and colocalized with the ERMCS SPLICS reporter sites (Fig. 4a), suggesting that the very close proximity with the ER is needed to promote the above-mentioned ultrastructural re-organization of mitochondria. We termed mitochondria with high ERMCS SPLICS^Hi^ and those are not associated with ERMCS as SPLICS^Lo^. Meanwhile, the SPLICS^Lo^ mitochondria with a higher ERMCS distance still form normal network-like morphology. However, OMM proteins (MFN2, DRP1, and phospho-DRP1^S616/637^), IMM organization proteins (OPA1, MICOS complex component MIC60 and MIC25), and mitochondrial electron transport chain (ETC) expression did not change in whole cell western blots (Extended Data Fig. 6a and b). In addition, expression of the ER sheet and tubule shaping proteins CLIMP63 (CKAP4) and RTN4 did not change (Extended Data Fig. 6c). These data suggest that the structural changes of ERMCS induction by fedratinib did not induce changes by altering levels of the canonical mitochondria dynamics or ER structure shaping proteins. However, we cannot rule out the role of membrane lipid remodeling that contributes to mitochondrial outer membrane and inner membrane structural changes.

**Fig. 4:**
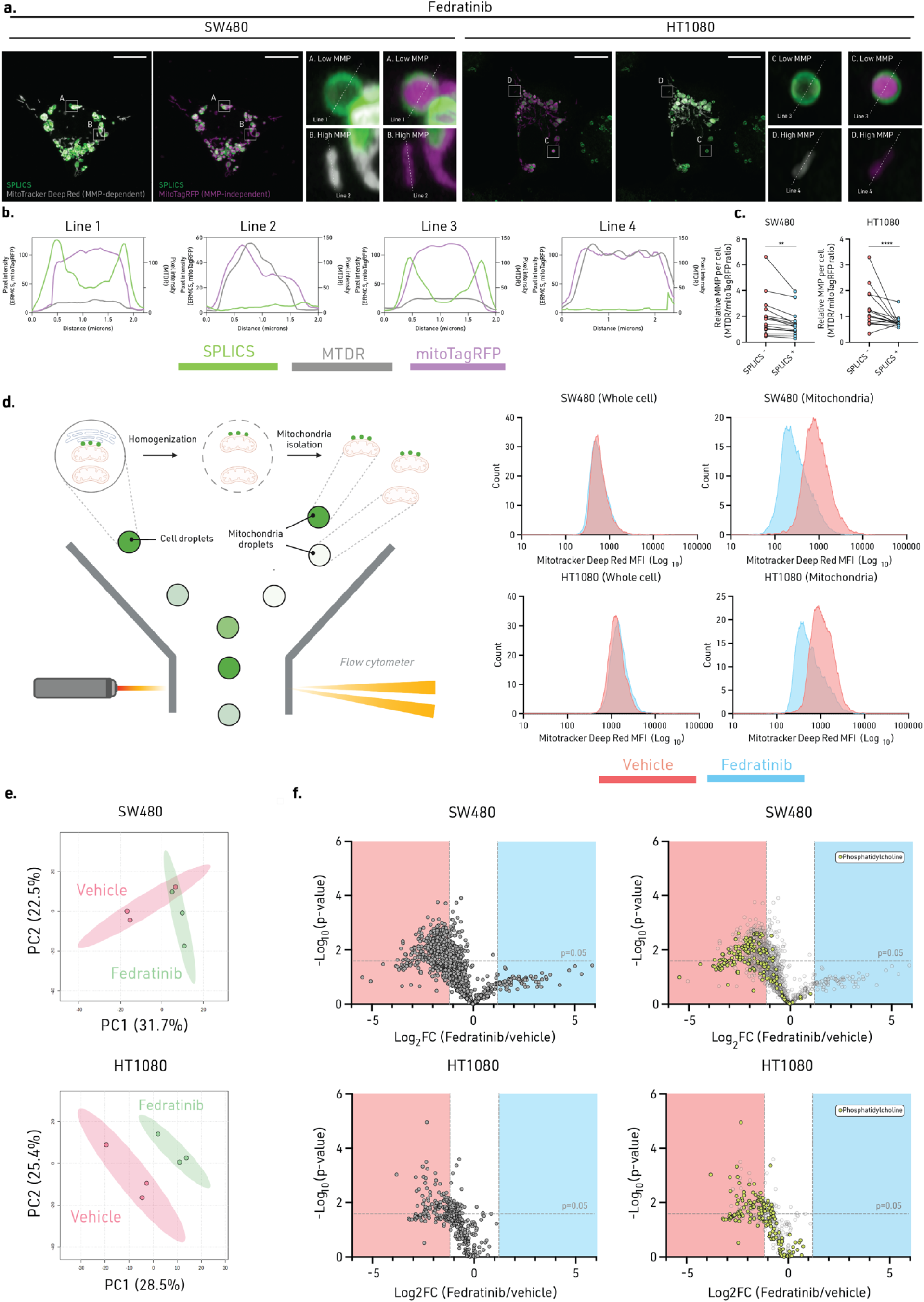
Metabolic and lipidomic changes associated with selective depolarization of ER-wrapped mitochondria. **a**, Confocal images and, **b**, line intensity profile of representative SPLICS^Hi^ and SPLICS^Lo^ mitochondria of mitochondria membrane potential (MitoTracker Deep Red (MTDR)) and total mitochondrial mass (mitoTagRFP) in cells treated with fedratinib for 24 hr. **c**, Ratio metric analyses of mitochondria membrane potential from SPLICS^Hi^ and SPLICS^Lo^ mitochondria in the same cell. Cells treated with fedratinib for 24 hr (n = 16; Scale bar: 1 μm). Each dot indicates one cell that was imaged. **d**, Diagram describing the FACS analysis of mitochondria, and mitochondria flow cytometry performed on whole cell (n = 10,000) or isolated mitochondrial (n = 100,000) fraction from cells treated with vehicle or fedratinib for 24 hr. log_10_-scaled MTDR mean fluorescence intensity (MFI) displayed on x-axis and count on the y-axis. **e**, Principal component analysis (PCA) of metabolomic analysis of cells treated with vehicle or fedratinib for 24 hr. **f**, Volcano plot of mitochondrial lipidome profiles in SW480 and HT1080 cells treated with fedratinib for 24 hr. **g**, Volcano plot of mitochondrial PtdCho species highlighted in green in cells treated with fedratinib for 24 hr. Experiments (**a**-**d**) were repeated three times. Experiment (**e**) was conducted in biological triplicates. Experiments (**f** and **g**) were conducted once with four biological replicates. Two-tailed paired t-test were performed on (E). **P ≤ 0.01, ***P ≤ 0.005. Statistical source data are provided in Source Data Fig.4.

To further characterize mitochondria function, we profiled mitochondrial mass, membrane potential (MMP/ΔΨm), oxygen consumption rate (OCR), extracellular acidification rate (ECAR), mitochondria reactive oxygen species (ROS), total cellular ROS, mitochondrial calcium response, and mitochondrial DNA (mtDNA) content. Again, we did not observe consistent differences between fedratinib versus control-treated cells in bulk measurements from whole cells (Extended Data Fig. 6c-g, i-k and Extended Data Fig. 7). From the seahorse OCR analysis, we analyzed basal respiration, maximal respiration, ATP-linked respiration, spare respiratory capacity (SRC), and proton leak. We observed a consistent decrease in maximal respiration ATP-linked respiration and SRC from both cell lines (Extended Data Fig. 7c, d, and e), suggesting ERMCS induction by fedratinib could limit adaptive respiratory changes. Since these mitochondrial phenotyping approaches report changes in the average behavior at a whole cell level changes, they could mask phenotypic differences since not all mitochondria are wrapped by ER in fedratinib-treated cells.

With these data in mind, we next employed single mitochondria functional profiling with fluorescence imaging, which revealed intra-cellular heterogeneity in mitochondria membrane potential. Mitochondria wrapped with ER (SPLICS^Hi^) have lower ΔΨm compared to those lacking ERMCS (SPLICS^Lo^) (Fig. 4a-c; Extended Data Fig. 8b). Orthogonal validation with single mitochondria flow cytometry further confirmed lower ΔΨm when ERMCS is induced that is absent at the whole cell level (Fig. 4d). Of note, we did not observe changes in mitochondrial ROS at the single mitochondrial or whole cell level (Extended Data Fig. 8a and c). Additionally, FACS analysis at the single mitochondrial level revealed higher mitochondrial calcium in SPLICS^Hi^ compared to SPLICS^Lo^ mitochondria (Extended Data. Fig. 6l).

These points notwithstanding, we investigated the impact of fedratinib on whole cell metabolism using metabolomics and lipidomics. Through dimensional reduction approaches to identify variability in metabolite and lipid profiles, we found that the metabolome is significantly altered upon induction of ERMCS (Fig.4e; Extended Data Fig. 9a-c; Supplementary Table 5). Indeed, the changes to the lipidome were even more profound when we performed lipidomics on mitochondria isolated from fedratinib-treated cells as we observed a dramatic decrease in mitochondrial lipids (Fig. 4f; Supplementary Table 6). Since loss of cardiolipin, a mitochondrial inner membrane lipid, or alteration in the phospholipid saturation state can directly affect cristae organization^30^. From our mitochondrial lipidomics that compared lipidome between fedratinib and vehicle treated cells, we identified a global decrease in lipids, particularly the structural phospholipid phosphatidylcholine (PtdCho) (Fig. 4f).

### Fedratinib induces subcellular proteomic changes specific to ER-enveloped mitochondria

To further explore mitochondrial heterogeneity within cells, we hypothesized that distinct mitochondrial populations, with varying degree of ER contact can be detected by single mitochondrial flow cytometry assay. Although flow cytometric analysis of ERMCS at the whole cell level revealed continuous distribution, analysis at the individual mitochondria level revealed two distinct population with high and low levels of SPLICS intensity. Specifically, 36.98% of mitochondria in SW480 cells and 46.30% in HT1080 cells are SPLICS^Hi^, whereas over 50% of the mitochondria exhibited low SPLICS signal (SPLICS^Lo^) (Fig. 5a). These data align with our microscopy data, which indicated only a subset of mitochondria were enveloped by the ER. Additionally, using forward scatter (FSC) as a proxy of particle size, we also validated that SPLICS^Hi^ mitochondria are larger than those not associated with the ER (Fig. 5b and c).

**Fig. 5:**
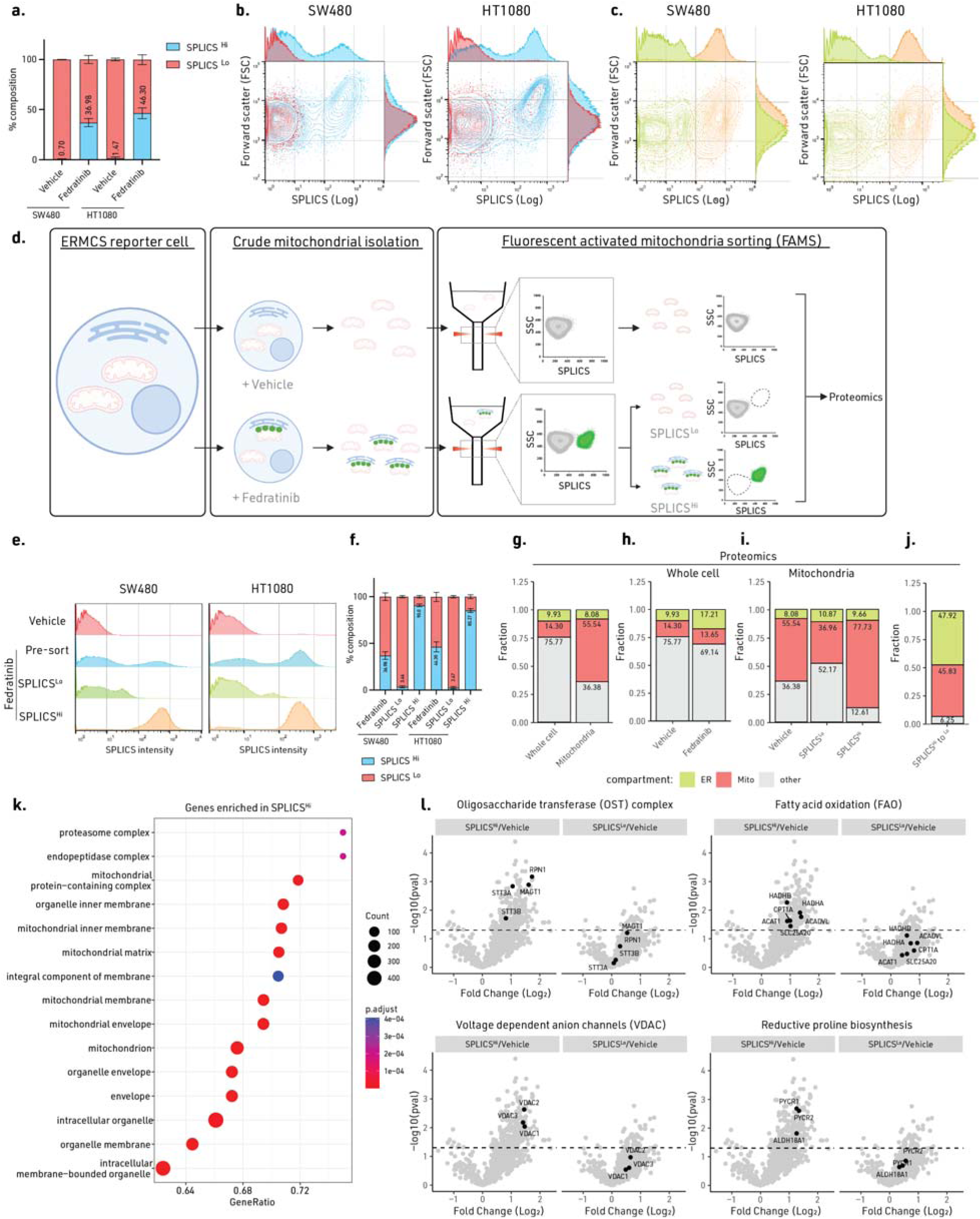
Fedratinib induces subcellular proteomic changes specific to ER-enveloped mitochondria. **a**, Distribution of SPLICS^Hi^ and SPLICS^Lo^ mitochondria in fedratinib and vehicle treated cells. **b**, Scatter plot of individual mitochondria from fedratinib and vehicle treated cells with x-axis as SPLICS MFI and y-axis as forward scatter (FSC). **c**, Scatter plot of sorted mitochondria from fedratinib and vehicle treated cells with x-axis as SPLICS MFI and y-axis as FSC. **d**, Schematic of mitochondrial flow cytometry and sorting for proteomics. **e**, Histograms of pre- and post-sorted mitochondria from fedratinib and vehicle treated cells. **f**, Distribution of SPLICS^Hi^ and SPLICS^Lo^ mitochondria in pre- and post-sorted, fedratinib-treated cells. **g**, Subcellular proteomic distribution of ER, mitochondria, and other compartmental proteins in vehicle-treated whole cell and sorted mitochondria. **h**, Subcellular proteomic distribution of ER, mitochondria, and other compartmental proteins in fedratinib-treated whole cell. **i**, Subcellular proteomic distribution of ER, mitochondria, and other compartmental proteins in sorted vehicle-treated mitochondria, SPLICS^Hi^ mitochondria, and SPLICS^Lo^ mitochondria. **j**, Subcellular proteomic distribution of ER, mitochondria, and other compartmental proteins comparing SPLICS^Hi^ to SPLICS^Lo^ mitochondria. **k**, GSEA analysis of SPLICS^Hi^ mitochondria. **l**, Volcano plot showing enrichment of proteins involved in: oligosaccharide transferase (OST) complex, fatty acid oxidation (FAO), voltage dependent anion channel (VDAC), and reductive proline biosynthesis. Experiments (**a**-**c**, **e**, **f**) were performed three times, experiments (**g**-**l**) were performed once with four biological replicates. Statistical source data are provided in Source Data Fig. 5.

To further identify mitochondrial level proteomic heterogeneity, we conducted fluorescence activated mitochondria sorting (FAMS) to isolate SPLICS^Hi^ and SPLICS^Lo^ mitochondria in fedratinib-treated cells relative to vehicle treated control (Fig. 5d)^31^. We successfully enriched for 90.8% of SPLICS^Hi^ in SW480 and 85.3% in HT1080 cells (Fig. 5e and f). We then conducted quantitative proteomics on the FAMS-sorted mitochondria from SW480 cells. Compared to the whole cell proteome, we enriched for mitochondrial protein by 3.8-fold (Fig. 5g). Fedratinib treatment increased ER localized protein around 2-fold, and all FAMS-sorted mitochondria have significant mitochondria enrichment (Fig. 5h and i). When comparing SPLICS^Hi^ to SPLICS^Lo^ mitochondria, we further enriched for ER proteins by approximately 4.8-fold (Fig. 5j), supporting higher degree of ER engagement with SPLICS^Hi^ mitochondria. Gene ontology (GO) analysis of SPLICS^Hi^ mitochondria identified significant enrichment of terms associated with mitochondria, membranes, and organelles (Fig. 5k).

When focusing on SPLICS^Hi^ mitochondria, we identified a strong enrichment of N-linked glycosylation machineries, including the catalytic subunits of the oligosaccharyltransferase (OST) complex STT3A and STT3B and other OST components such as RPN1 and MAGT1 (Fig. 5l). Additionally, mitochondrial proteins selectively enriched in SPLICS^Hi^ mitochondria, included voltage dependent anion channels (VDAC) VDAC1, 2, and 3, which are known ERMCS tethers. Interestingly, we also identified enrichment of enzymes involved in reductive proline biosynthesis genes ALDH18A1/P5CS, PYCR1, and PYCR2, as well as fatty acid oxidation (FAO) proteins CPT1A, ACADVL, HADHA/B, ACAT1, and SLC25A20. This suggests that mitochondria in contact with ER could be involved in distinct metabolic processes compared to those that are not interacting with the ER (Figure 5l). Through our single mitochondrial level analyses, we have identified unique proteome associated with mitochondria in physical contact with the ER.

### ETC Complex III function is necessary for ERMCS formation

Based on the drastic mitochondrial structural and less pronounced but significant metabolic changes associated with ERMCS induction, we hypothesized that an optimal metabolic state would be required to sustain metabolic demands for ERMCS formation. To test this model, we employed a variety of mitochondrial perturbations, and found that hypoxia reduced ERMCS (Fig. 6a). In addition to directly inhibiting oxygen dependent-reactions^32^, hypoxia also rewires cellular metabolism through hypoxia inducible factors (HIFs). In contrast to hypoxia, HIF induction with the PDH inhibitor FG-4592 did not similarly attenuate fedratinib-induced ERMCS, indicating that the hypoxia effect is independent of HIF activation but oxygen dependent (Fig. 6a, extended figure 10a).

**Fig. 6:**
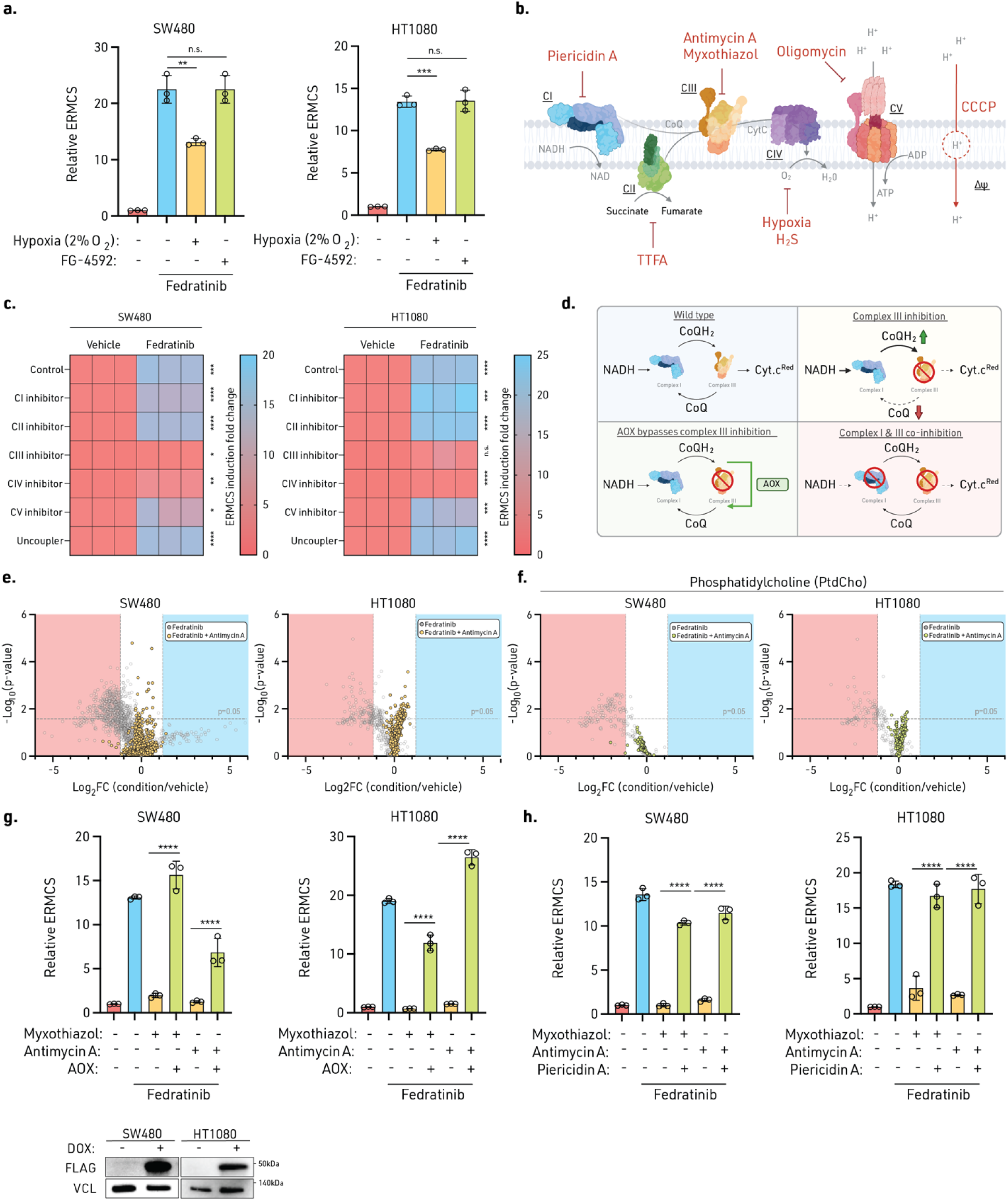
ETC Complex III function is necessary for ERMCS formation. **a**, Relative ERMCS (right) in cells incubated in hypoxia (2% O_2_) or treated with a PHD inhibitor FG-4592 (50 μM) for 48 hr, and then treated with fedratinib for 24 hr. **b**, Diagram of mitochondria electron transport chain (ETC) and specific mitochondrial complex inhibitors (CI: piericidin A (20 nM), CII: TTFA (100 μM), CIII (antimycin A: 100 nM, myxothiazol: 50 nM), CIV (Hypoxia: 2% O_2_, H_2_S: 250 ppm)). **c**, Relative ERMCS (right) in cells treated with specific ETC inhibitors for 24 hrs, and then treated with vehicle or fedratinib for 24 hr. **d**, Diagram illustrating the regulation of CoQ redox state following complex III inhibition, expression of alternative oxidase (AOX) under complex III inhibition, and complex I/III dual inhibition. **e**, Volcano plot of mitochondrial lipidome profiles comparing fedratinib (grey circle) and fedratinib plus antimycin A combination treatment (yellow dot). **f**, Volcano plot of mitochondrial PtdCho species in fedratinib (grey circle) and fedratinib plus antimycin A combination treatment (yellow dot). **g**, Relative ERMCS in AOX expressed cell treated with complex III inhibition with myxothiazol (50 nM) or antimycin A (100 nM) for 24 hr, and then treated with vehicle or fedratinib for 24 hr. AOX expression was validated with immunoblotting (below). **h**, Relative ERMCS in cells treated with complex I and III inhibitors Piericidin A (20 nM) and myxothiazol (50 nM), or antimycin A (100 nM) for 24 hr, and then treated with vehicle or fedratinib for 24 hr. Experiments (**a**, **c**, **g**, **h**) were performed three times. Experiments (**e** and **f**) were performed once with four biological replicates. Significance was calculated with an unpaired two-tailed t-test. n.s. = not significant, **P ≤ 0.01, ***P ≤ 0.001, ****P ≤ 0.001. Data are mean ± SD. Statistical source data are provided in Source Data Fig. 6. Immunoblot source data are provided in Supplemental Figure 1.

Since mitochondria are the largest oxygen consumers within the cell, we individually inhibited the electron transport chain (ETC) complexes I-V, or depolarized membrane potential, and then asked whether a specific ETC component is required for ERMCS induction (Fig. 6b). Only complex III and IV inhibition suppressed the formation of ERMCS induced by fedratinib (Fig. 6c). We also report that CIII inhibition with Antimycin A reversed mitochondrial PtdCho and lipid depletion from fedratinib treatment (Fig 6e and f), connecting consistent regulatory patterns between ERMCS and mitochondrial lipid regulation.

Unlike ETC inhibitors that restrict oxygen consumption and generate mitochondrial ROS, inhibiting complex III or IV suppresses the ability to carry out CoQ/ubiquinol oxidation. Inhibition of complex III and IV leads to the buildup of CoQH_2_, which leads to a reductively shifted CoQH_2_/CoQ pool, which has been associated with reverse electron transport (RET) at complex I and II, generating ROS and reducing fumarate to succinate^33^, respectively.

We found that neither mitochondrial-specific ROS inducer mito-paraquat (mitoPQ) nor a complex I inhibitor that increases oxidized CoQ pool suppressed ERMCS formation (Extended Data Fig. 10b). Moreover, blocking electron leak and superoxide generation at complex III Q_0_ site with S3QEL-2 did not rescue antimycin dependent ERMCS suppression (Extended Data Fig. 6h). Therefore, we hypothesized that complex III inhibition could suppress ERMCS formation from accumulation of reduced CoQH_2_, but not the generation of ROS. To relieve CoQH_2_ buildup, we genetically expressed a *Ciona intestinalis* alternative oxidase (AOX)^33^, which oxidizes CoQH_2_ to CoQ in an antimycin/myxothiazol-insensitive manner (Fig. 6d). If CoQH_2_ accumulation drives ERMCS suppression, AOX expression should restore ERMCS induction by maintaining an oxidized CoQ pool. Notably, cells expressing AOX retain fedratinib-induced ERMCS when simultaneously subjected to complex III inhibitors (Fig. 6g; Extended Data Fig. 10c). This regulation is not limited to fedratinib as thapsigargin induced ERMCS are also regulated by similar mechanisms (Extended Data Fig. 10d and e). Surprisingly, fedratinib did not directly alter the CoQH_2_/CoQ redox ratio or CoQ_10_ abundance at the whole cell or mitochondria level when measured by mass spectrometry and high performance liquid chromatography (HPLC) (Extended Data Fig. 10h and i). This suggest that perhaps maintaining the ability for complex III to carry out CoQ oxidation, rather than the total levels, is crucial for ERMCS induction. Through confocal microscopy, we also showed that ER-wrapped mitochondrial are enriched with CoQ biosynthetic enzyme COQ9, indicating a potential role in local CoQ channeling capabilities (Extended Data Fig. 10j).

In addition to expressing AOX, we also utilized an orthogonal method to alter electron flow to complex III. Under standard conditions, complex I contributes the majority of electrons to the ETC via NADH. Co-inhibiting complex I and complex III limits CoQH_2_ generation, thereby decreasing the CoQH_2_/CoQ ratio. Indeed, inhibition of complex I and III restored ERMCS inducibility (Fig. 6h). We attempted to rescue the impact of complex IV inhibitors, H_2_S and hypoxia, by similarly inhibiting complex I. However, piericidin A was unable to restore ERMCS levels (Extended Data Fig. 10f and g). This may be attributed to the dominant role of SQOR-mediated CoQ reduction as the main electron donor under H_2_S conditions, as well as the inhibition of NADH-dependent electron flux into the ETC during hypoxia.

These findings support that complex III-dependent functions such as CoQ oxidation could influence the dynamic induction of ERMCS formation. More importantly, this regulation is not limited to fedratinib induced ERMCS, as ERMCS induced by thapsigargin also responds to methods that bypass Complex III or CoQ oxidation state manipulation. Consequently, we can conclude that the Complex III regulated CoQ oxidation could act as a reliable indicator of ERMCS dynamics and is a novel metabolic pre-requisite for ERMCS induction.

## Discussion

Inter-organellar contact sites mediate local signaling events, whole cell metabolism and tissue physiology. The dysregulation of ERMCS has been implicated in diseases including cancer, fragile X syndrome^34^, and Friedrich’s ataxia^13^. Despite their critical role in maintaining metabolic homeostasis, the mechanisms governing the dynamics of ERMCS remain elusive, at least in part due to the lack of potent inducers with temporal control. We identified fedratinib, an FDA-approved drug for myelodysplastic syndrome that robustly induced the formation of ERMCS across diverse cell lines. We demonstrated that the relevant molecular target of fedratinib was the epigenetic regulator BRD4, and inhibition initiated an activating gene expression program that coordinated with mediator complex to induce ERMCS. Using iterative EM approaches, starting with TEM, CLEM, and then cryo-preserved FIBSEM, we identified that ERMCS exhibited novel ultrastructural changes in response to fedratinib. These changes included a tight and complete envelopment of the entire mitochondrial membrane at a subset of mitochondria.This is similar to ER-mitochondrial interactions observed following infection with human cytomegalovirus and SARS-CoV-2, metastatic melanoma cells, and fasted hepatocytes in liver^35–37^.

Despite these structural alterations, no consistent changes were observed in various ER and mitochondrial functions at the whole-cell level. This is remarkable, given that approximately 30% of mitochondria exhibited massive structural alternations, indicating that “normal” mitochondria may compensate to support cell homeostasis when other mitochondria are dysfunctional. To dissect mitochondrial level heterogeneity, we optimized a single mitochondrial sorting pipeline to distinguish different mitochondrial populations. As we observed from our single mitochondria proteomics, mitochondria with abundant contact sites with the ER may potentially be dedicated to support specific metabolic pathways, including proline biosynthesis or beta oxidation. These results underscore the significance of considering mitochondrial heterogeneity in the study of ERMCS. Moreover, through probing ETC function, we demonstrated that ERMCS formation is dependent on the activity of mitochondria complex III/IV. Methods bypassing their function restored ERMCS abundance, suggesting that non-affected mitochondria adapt or persist when complex III function is suppressed, whereas the affected mitochondria are targeted for ER engulfment.

Previous studies of BRD4 predominantly focused on its ability to act as a positive regulator of transcription^38^. In contrast, our findings demonstrate that de-repression of its target gene expression is required for ERMCS activity. Inhibition of BRD4 activates autophagy-related genes and corrects for mitochondrial disease phenotype by increasing mitochondrial biogenesis^28,39^. However, the direct transcriptional target(s) of BRD4 that regulate ERMCS are not clear. Through transcriptomics we identified activation of the ISR in both cells with high basal ERMCS and following fedratinib treatment. ISR activation relies on inducing PERK, HRI, GCN2, and PKR^40^. Recent work has demonstrated that mitochondrial stress and sublethal activation of mitochondrial outer membrane permeabilization (MOMP) can release cytochrome c and subsequently activate ISR^41^. Interestingly, the loss of cristae at ERMCS resembles the effect induced by MOMP activation^42^. Moreover, the dynamic nature of ERMCS is involved in apoptosis regulation and sensitivity^43,44^. Since the UPR can activate the ISR, we did extensive comparison of fedratinib to thapsigargin. We did not observe UPR activation after fedratinib treatment nor changes in UPR targets (IRE1α, PERK, ERO1L, TMX1) that have been associated with ERMCS induction^45–47^. However, direct activation of the ISR kinase PKR induced ERMCS, while inhibiting PKR suppressed fedratinib induced ERMCS. Interestingly, ISR inhibitor ISRIB decreased fedratinib-dependent ATF4 upregulation but failed to suppress ERMCS. These findings reveal compelling connections between ISR signaling and ERMCS regulation, underscoring the complex interactions among eIF2α kinases and distinct cellular effects from early versus latent ISR activation. Discerning the relationship between ISR and ERMCS warrants further study.

The ratio of oxidized CoQ to reduced CoQH_2_ plays a critical signaling role that reflects cellular metabolic states. While recognized as an electron carrier in the ETC ^48^, CoQ has emerged as a multifunctional lipid involved in ferroptosis^49,50^, pyrimidine biosynthesis^51^, H_2_S detoxification^52^, glycerol phosphate shuttle, and proline catabolism^53^. Accumulation of CoQH_2_ under hypoxia or sulfide exposure is essential to maintain tissue homeostasis by utilizing fumarate as a terminal electron acceptor instead of oxygen^33,52^. In physiological or pathological hypoxia, tissues rewire metabolism via the canonical HIF oxygen sensing pathway and reorganize organellar networks to maintain metabolism and other aspects of cellular homeostasis. Indeed, we demonstrated that ETC complex III and IV could sustain a critical function needed for establishing ERMCS as inhibiting complex III and IV inhibited ERMCS. Utilizing genetic methods to bypass complex III inhibition via ectopic expression of alternative oxidase (AOX) restored ERMCS abundance under complex III inhibition. Surprisingly, limiting electron entry from complex I reversed the inhibitory response of antimycin or myxothiazol and re-activated ERMCS induction. Based on the AOX and complex I/III co-inhibition results, we speculate that there could potentially be a CoQ redox sensing mechanism as both rescue approaches should prevent CoQH_2_ accumulation under complex III inhibition alone. Further studies to profile CoQ oxidation state on the subcellular level and spatial distribution will be critical for drawing definitive conclusions with the inconsistent change in CoQ redox state and CoQ abundance. Thus, we provide evidence to support that modulation of oxygen tension re-organizes the subcellular organellar landscape and Complex III function could potentially be an endogenous metabolic sensor for ERMCS formation. Importantly, while we made this discovery with fedratinib, we also illustrated that other ERMCS inducers, like the ER stressor thapsigargin, require the same molecular and metabolic mechanisms for ERMCS. This evidence suggests the mechanisms described here are broadly applicable to other ERMCS regulatory modalities.

In conclusion, our study underscores the intricate interplay between molecular regulators and metabolic cues that dictate the formation and functionality of ERMCS. Moreover, we characterize a tool compound that can temporally and reversibly modulate ERMCS laying the foundation for potential therapeutics targeting inter-organellar communication in diverse disease contexts.

## Materials and Methods

### Cell culture

HT1080, SW480 HCT116, DLD1, SW480, HeLa, U2OS, A459, PA-TU8902, K562, Jurkat, B16, KPC7940 cells were maintained at 37°C in 5% CO_2_ and 21% O_2_. HT1080, SW480 HCT116, DLD1, SW480, HeLa, U2OS, A459, PA-TU8902, B16, and KPC7940 cells were cultured in Dulbecco’s Modified Eagle Medium (DMEM) supplemented with 10% FBS and 1% antibiotic/antimycotic, whereas Jurkat and K562 cells were cultured in Roswell Park Memorial Institute (RPMI) supplemented with 10% FBS and 1% antibiotic/antimycotic.

### Generation of SPLICS stable cell lines

The SPLICS reporter is composed of a split GFP_1-10_ barrel localized to the mitochondrial outer membrane, a P2A self-cleavage signal for equimolar expression, and the remaining beta_11_ fragment targeted to the ER membrane. Upon ERMCS formation within the 8-10 nm distance, the split-GFP reporter forms intact GFP. Doxycycline-inducible SPLICS reporter stable cell lines were generated via a three-plasmid PiggyBac transposase system. Cells were co-transfected with the plasmids using Lipofectamine 2000 transfection reagent, then the next day selected with 2 mg/ml of G418 (Geneticin) for 7 days. After selection, 100 ng/ml of doxycycline were used to induce reporter expression, and GFP positive cells were sorted on Bigfoot Spectral Cell Sorter (Invitrogen) to isolate GFP positive clones. Clones were cultured and screened for accurate localization of SPLICS reporter to mitochondria and ER via live cell imaging and for normal mitochondrial oxygen consumption rate. For ERMCS analysis with SPLICS, cells will be induced with 100 ng/ml of doxycycline for minimally 24 hr before treatment or analysis.

### Generation of genetically modified cell lines

sgRNAs (oligonucleotide sequences are indicated in Supplementary Table 5) were ligated into BsmBI-linearized lentiCRISPR-v2 (for CRISPR KO) or pLV hU6-sgRNA hUbC-dCas9-KRAB-T2a-Puro (pLV hU6-sgRNA hUbC-dCas9-KRAB-T2a-Puro was a gift from Charles Gersbach (Addgene plasmid # 71236; http://n2t.net/addgene:71236; RRID:Addgene_71236)) with T4 ligase (NEB). AOX overexpression plasmid was achieved with pCW57.1_AOX-FLAG (pCW57.1_AOX-FLAG was a gift from David Sabatini & Jessica Spinelli (Addgene plasmid # 177984; http://n2t.net/addgene:177984; RRID:Addgene_177984)). MitoTagRFP plasmid was acquired from Addgene (pclbw-mitoTagRFP was a gift from David Chan (Addgene plasmid # 58425; http://n2t.net/addgene:58425; RRID:Addgene_58425). BFP and BFP-ER-mitochondria tether were generated via Vectorbuilder (pLV[Exp]-Bsd-CMV>{new mTagBFP2} and pLV[Exp]-Bsd-CMV>{new mTagBFP2 ER mito tether}). Lentiviral vectors expressing sgRNAs were transfected into HEK293T cells with lentiviral packaging vectors CMV VSV-G and psPAX2 using XtremeGene 9 transfection reagent (Roche). After 24 h, media was aspirated and replaced by fresh media. The virus-containing supernatant was collected 48 h after transfection and filtered using a 0.45 mm filter to eliminate cells. Cells to be transduced were plated in 6-well tissue culture plates and infected in media containing virus and 8 μg/mL of polybrene. Cells were spin infected by centrifugation at 1,100 g for 1.5 h. After transduction, media was changed, and cells were selected with puromycin (for sgRNA lentiviral vector) for 72 hrs.

### siRNA mediated knockdown

siRNAs for BRD4 were transfected using a reverse transfection protocol and Lipofectamine RNAiMAX. siRNA was purchased from Horizon Discovery with the following information: siGENOME Human BRD4 (23476) siRNA - SMARTpool, 5 nmol.

### Western Blots

Whole-cell lysate preparations were described previously (Anderson et al., 2013). Whole cell lysates were prepared from cell lines by RIPA buffer. Homogenates were incubated in RIPA buffer for 15 min on ice followed by 13,000 rpm centrifugation for 15 min. Supernatants were transferred to a new tube and mixed with 5× Laemmli buffer and boiled for 5 min. Lysates containing 30–40 μg of protein per well were separated by SDS-PAGE, transferred onto nitrocellulose membranes, and immunoblotted overnight at 4°C with indicated antibodies. All the primary antibodies were used at a dilution of 1:1000. HRP-conjugated secondary antibodies used were anti-rabbit and anti-mouse at a dilution of 1: 2000 and immunoblots were developed using the iBright Imaging System and iBright Analysis Software (Thermofisher).

| Antigen | Company | Catalog number |
| --- | --- | --- |
| Vinculin (clone: E1E9V) | Cell Signaling Technology | 13901 |
| MFN2 (clone: D2D10) | Cell Signaling Technology | 9482 |
| DRP1 (clone: D6C7) | Cell Signaling Technology | 8570 |
| pS616-DRP1 | Cell Signaling Technology | 3455 |
| pS637-DRP1 | Cell Signaling Technology | 4867 |
| Mitofilin/MIC60/IMMT | Proteintech | 10179-1-AP |
| MIC25/CHCHD6 | Proteintech | 20639-1-AP |
| RTN4/NOGO | Proteintech | 10740-1-AP |
| COQ9 | Proteintech | 14874-1-AP |
| CLIMP63/CKAP4 (clone: G1/296) | Enzo Life Science | ENZ-ABS669-0100 |
| OPA1 (clone: D6U6N) | Cell Signaling Technology | 80471 |
| ATF4 (clone: E4Q4E) | Cell Signaling Technology | 97038 |
| BiP/HSPA5 (clone: C50B12) | Cell Signaling Technology | 3177 |
| CHOP/DDIT3 (clone: L63F7) | Cell Signaling Technology | 2895 |
| BRD4 (clone: E2A7X) | Cell Signaling Technology | 13440 |
| BRD4 (For fractionation) | Bethyl | BL-149-2H5 |
| c-MYC (clone: E5Q6W) | Cell Signaling Technology | 18583 |
| STAT3 (clone: 123H6) | Cell Signaling Technology | 9139 |
| pY705-STAT3 (clone: D3A7) | Cell Signaling Technology | 9145 |
| IP3R-I/II/III (clone: B-2) | Santa Cruz Biotechnology | SC-377518 |
| TOM20 (9D8T4N) | Cell Signaling Technology | 42406 |
| CCPG1 (E3C5G) | Cell Signaling Technology | 80158 |
| PERK (C33E10) | Cell Signaling Technology | 3192 |
| IRE1α (14C10) | Cell Signaling Technology | 3294 |
| ATF6 (D4Z8V) | Cell Signaling Technology | 65880 |
| XBP1s (E9V3E) | Cell Signaling Technology | 40435 |
| LC3B (D11) XP | Cell Signaling Technology | 3868 |
| HIF1α (D2U3T) | Cell Signaling Technology | 14179 |
| JAK2 (D2E12) XP | Cell Signaling Technology | 3230 |

### CRISPR Screen

For the epigenetics library screen in SW480 reporter cells, the human sgRNA library was described in this paper^54^ was used. The titer of lentiviral supernatants was determined by infecting targets cells at several amounts of virus in the presence of polybrene (10 ug/mL), counting the number of drug resistant infected cells after 3 days of selection. 40 million target cells were infected at an MOI of ∼0.5 and selected with puromycin (1 ug/mL) 72 h after infection. An initial pool of 30 million cells was harvested for genomic DNA extraction. The remaining cells were cultured for 6 days, treated with fedratinib for 24 hr, then 25 million cells were sorted from the top and bottom 10% of SPLICS/mitoTagRFP fluorescent ratio on day 8. After which cells were harvested for genomic DNA extraction. sgRNA inserts were PCR amplified (primers listed in Supplementary Table 7, purified and sequenced on a NextSeq (Illumina). Sequencing reads were mapped and the abundance of each sgRNA was tallied. Gene score is defined as the median log2 fold change in the abundance between the initial and final population of all sgRNAs targeting that gene. Full result of the screen with sequencing primers can be found in the supplemental data.

### CRISPR screen analysis

sgRNA read counts were generated by aligning sequencing reads to the sgRNA library sequences, with only exact matches allowed. sgRNA abundances were calculated for both sorted populations by first adding a pseudocount of 1, then dividing by the total number of reads in each given sample. For each sgRNA, a differential score was calculated as the log2 fold-change in sgRNA abundance between the GFP-high (top 10%) and GFP-low (bottom 10%) cell populations. Gene level differential scores were generated by taking the mean score of all sgRNAs targeting the same gene. The set of differential gene scores was then standardized by subtracting the mean from each gene score then dividing by the standard deviation (Z-score).

### Gene set enrichment analysis (GSEA) for CRISPR screen and proteomics

Gene set enrichment analysis was performed on the set of standardized differential gene scores using the clusterProfiler (v4.2.2) package in R. The gseGO() function was called on the data using all ontologies, min and max GSSsizes set to 10 and 500, respectively, FDR as the p-adjustment method, and an FDR cutoff of 0.01.

### Imaging based drug repurposing screen

384-well, black, optically clear flat-bottom, tissue-culture treated plates (Perkin Elmer cat# 6057302) were coated with 0.1mg/ml of poly-D-lysine hydrobromide (Sigma-Aldrich cat# P6407-5MG) at 37°C for 1hr. Then each plate is washed with molecular grade sterile water once, and then dried at room temperature for 1 hr. Each plate was then dry-spotted with 50nL of 2mM drug solution in DMSO using a Beckman Echo 650 acoustic liquid handler. In each plate, we have 2 columns of vehicle control (Dimethylsulfoxide/DMSO) and 2 columns of Thapsigargin (50 nM) as positive control for inducing endoplasmic reticulum-mitochondrial contact sites. With the plates coated and dry-spotted with drugs, 5000 SW480 colorectal cancer cells were plated in each well with complete DMEM media with a final drug concentration at 2μM. After incubation with the drug for 24 hr, cells were washed with PBS 3 times then fixed with 4% paraformaldehyde (PFA) in 1x PBS (Thermofisher, cat#043368.9M) at room temperature for 15 min in the dark. PFA was removed with 3 PBS washes and the fixed cells were staining with Hoechst 33342 (Invitrogen, cat# H3570, final concentration: 5 μg/ml) and HCS CellMask Deep Red Stain (Invitrogen, cat# H32721, final concentration: 1 μg/ml) for 1 hr at 37°C for 1hr. After staining, the cells were washed with PBS 3 times, and were left in PBS in the dark at 4°C until imaging. Plates were imaged on a Yokogawa CellVoyager 8000 using a 40X/1.0NA water immersion objective lens and nine fields per well were imaged across three channels to visualize Hoechst, GFP, Mitotag RFP and HCS CellMask Deep Red. Cellpose was used to identify nuclei and a custom CellProfiler pipeline was used to delineate the whole cell, mitochondria, GFP spots and per-cell feature extraction was performed. Endoplasmic reticulum–mitochondria contact sites (ERMCS) were identified by counting GFP spots within the cell. A XGBoost machine learning model was trained on centered/scaled cell-level features trained against the negative control (DMSO) and thapsigargin treated positive control wells to enable hit picking. XGBoost models were trained on the screening data controls from the entire screen with an 80%/20% train/test split. Model performance was excellent with a R2>0.95. Z-prime factors were calculated for each plate and plates below 0.5 were rejected and re-screened. High correlation was found between the XGBoost score and the GFP spot count per-cell, however, the XGBoost score exhibited significantly higher separation between the controls and led to higher Z-primes.

A liberal hit threshold of 3 standard deviations above the mean of the negative control (μ+3σ) was used to maximize hit detection despite an increase in the false positive rate. False positives were eliminated in dose-response confirmation where 64 compounds were tested in 10-point:2-fold dilution from 10uM to 20nM in triplicate wells. Of the 64 compounds tested, 33 were found to be dose-responsive (52% confirmation rate). This is typical for a bioactive compound library where the compound potencies range from sub-nanomolar to micromolar range and, often, potent hits are obscured by cytotoxicity when screening at low micromolar concentrations. Hits were cherry picked and confirmed with triplicate 10-point/2-fold dilution and EC50 values were fit. Images and details of the screen were deposited to Zenodo and are publicly available in the repository as part of this record: DOI: <u>10.5281/zenodo.13963526</u>.

### Oxygen Consumption Rate (Seahorse)

Cells were seeded at 2 × 104 cells/well in 80 μl/well of normal growth media (DMEM with 25 mM Glucose and 2 mM Glutamine) in an Agilent XF96 V3 PS Cell Culture Microplate (#101085-004). To achieve an even distribution of cells within wells, plates were incubated on the bench top at room temperature for 1 h before incubating at 37 °C, 5% CO2 overnight. To hydrate the XF96 FluxPak (#102416-100), 200 μL/well of sterile water was added and the entire cartridge was incubated at 37 °C, no CO2 overnight. The following day, 1 h prior to running the assay, 60 μL/well of growth media was removed from the cell culture plate, and cells were washed twice with 200 μL/well of assay medium (XF DMEM Base Medium, pH 7.4 (#103575-100) containing 25 mM glucose (#103577-100) and 2 mM glutamine (#103579-100)). After washing, 160 μL/well of assay medium was added to the cell culture plate for a final volume of 180 μL/well. Cells were then incubated at 37 °C, without CO2 until analysis. One hour prior to the assay, water from the FluxPak hydration was exchanged for 200 μL/well of XF Calibrant (#100840-000) and the cartridge was returned at 37 °C, without CO2 until analysis. Oligomycin (100 μM), FCCP (100 μM), and Rotenone/Antimycin (50 μM) from the XF Cell Mito Stress Test Kit (#103015-100) were re-constituted in assay medium to make the indicated stock concentrations. Twenty microliters of Oligomycin was loaded into Port A for each well of the FluxPak, 22 μL of FCCP into Port B, and 25 μL of Rotenone/Antimycin into Port C. Port D was left empty. The final FCCP concentration was optimized to achieve maximal respiration in each condition. The Mito Stress Test was conducted on an XF96 Extracellular Flux Analyzer and OCR was analyzed using Wave 2.6 software. Following the assay, OCR was normalized to cell number utilizing Agilent Cytation5 live cell imaging with Gen5 software.

### MT-DNA Quantification

Cells were pelleted, washed 1X in PBS, and then lysed in buffer (25 mM NaOH, 0.2 mM EDTA) for 15 minutes at 95°C. Lysis was neutralized with buffer (40 mM Tris-HCl) and centrifuged at 16,000 x g for 10 minutes at 4°C. Supernatant containing MT-DNA was quantified on with nanodrop. Primers amplifying the MT-DNA marker D-Loop and the nuclear DNA marker β-Globin were used and the relative MT-DNA to nuclear DNA was quantified in each sample^55^.

### Mitochondria isolation

SW480 and HT1080 cells were scraped and pelleted at 1000 x g for 5 minutes. Cell pellets were washed in 1X PBS and pelleted at 1000 x g for 5 minutes. Cells were re-suspended in isolation buffer (200 mM sucrose, 10 mM Tris HCl, 1 mM EGTA/Tris, pH 7.4 (adjusted with 1M HEPES)), transferred into a 27-gauge needle and homogenized by 10 repeated resuspensions. Homogenates were transferred into tubes and spun at 600 x g for 10 minutes at 4°C. Supernatants were moved to a new tube and centrifugation was repeated. Then, supernatants were moved to a new tube and pelleted at 7,000 x g for 10 minutes at 4°C. Pellets were washed in isolation buffer and centrifugation was repeated two more times. For biochemical analyses, mitochondria were aliquoted in 50 μg pellets.

### Flow cytometry for cells

HT1080 and SW480 cells were seeded 48 hr to 72 hr prior to the experiment at 50K and 75K cells density to achieve final 70% confluency by the time of analysis. For analyzing SPLICS and mitoTagRFP intensity, cells were washed, trypsinized, and filtered through a cell strainer prior to analyzing. For MitoSOX Deep Red(Dojindo, cat#: MT14-12), MitoTracker Green (Invitrogen, cat#: M7514), Tetramethylrhodamine, Ethyl Ester, Perchlorate/TMRE (Invitrogen, cat#: T669), and Rhod-2, AM (Invitrogen, cat#: R1244) experiments, cells were stained with 1 μM of MitoSOX Deep Red, 100 nM of MitoTracker Green, 500 nM of TMRE, 10μM of DAPRed (stained for 30 minutes, and then gets replaced in regular media) and 5μM of Rhod-2 for 30 min 37 °C, washed twice with warm PBS, and proceeded with the same processing procedure as unstained cells. Fluorescence activated cell sorting (FACS) was performed on Bigfoot Spectral Cell Sorter (Invitrogen), in which the fluorescence intensity of minimum 10K individual cells was quantified in their specific channel. Data were analyzed using FlowJo software (TreeStar).

### Flow cytometry for mitochondria

HT1080 and SW480 cells were seeded at 1.5 million and 3 million in a 15cm plate to achieve final 70% confluency by the time of analysis. Upon 24 hr of treatment with Fedratinib cells were stained with MitoTracker Deep Red or MitoSOX Deep Red. Upon staining, 10% of whole cell fraction was set aside and then mitochondria were isolated with the protocol mentioned. Briefly, cells were fixed at room temperature for 2 min with 2% PFA, and then washed in mitochondria isolation buffer. FACS was performed on Bigfoot Spectral Cell Sorter (Invitrogen), in which the fluorescence intensity of minimum 10K individual cells or 100K mitochondria was quantified in their specific channel. Data were analyzed using FlowJo software (TreeStar).

### Fluorescence activated mitochondria sorting (FAMS)

SW480 and HT1080 SPLICS/mitoTagRFP-expressing cells are plated at 3e^6^ and 6e^6^ respectively in a 150-mm plate 48 hours before sorting with doxycycline treatment to induce SPLICS expression. Cells were treated with fedratinib or vehicle (DMSO) control overnight. The next morning, cells were stained with MitoBrilliant 646 (Tocris #7700) at 200 nM for 1 hour prior to mitochondrial isolation. Mitochondria isolations were conducted as previously indicated. FAMS was conducted on the Bigfoot Spectral Cell Sorter (Invitrogen) with 70-micron nozzle. The fluidics were flushed with deionized water for 5 minutes followed by equilibration with mitochondrial isolation buffer for another 5 minutes. Polystyrene beads (Spherotech) with defined sizes (0.2 μm, 0.5 μm, 1 μm, and 2 μm) from were used to determine gating size of mitochondria particles. After gating for size, mitochondria populations were determined using mitoTagRFP and MitoBrilliant 646 double positive gates. The populations then were analyzed based on GFP signal. SPLICS^Lo^ mitochondria are determined based on vehicle control, and SPLICS^Hi^ mitochondria are separated based on the formation of distinct GFP population in fedratinib-treated cells. For proteomics, contain 60 million mitochondria were sorted in each condition and four replicates were collected.

### Live cell confocal imaging

35mm glass bottom plates (Ibidi) were coated with 0.1mg/ml of poly-D-Lysine (Sigma) for 1 hour at 37°C, washed once with sterile water, dry in the biosafety cabinet for 2 hours, and then complete media was added on to equilibrate for 20 min in the incubator. 50K of HT1080 and 100K of SW480 cells were then plated to achieve final 60-70% confluency by the time of imaging. Cells with HaloTag-Sec61 were stained with 200nM of JaneliaFluor 646 (JF646, Promega) for 30min at 37°C, and then washed once with warm PBS. Cells were counterstained with 2000x dilution of Hoechst 33342 (Thermofisher) for 15min at 37°C. FluoroBrite DMEM was used as the media for live cell imaging. The cells were imaged using Zeiss LSM 980 Airyscan 2 microscope and detector equipped with a 63x oil objective with a 1.4 numerical aperture. CO_2_ and humidity was equilibrated for at least 30 min before imaging. Post-processing was done with Zen 3.4 (Blue edition).

### Immunofluorescence (IF)

Number 1.5 round 10-mm glass coverslips were coated with 0.1mg/ml of poly-D-Lysine (Sigma) for 1 hour at 37°C, washed once with sterile water, dry in the biosafety cabinet for 2 hours, and then complete media was added on to equilibrate for 20 min in the incubator. 50K of HT1080 and 100K of SW480 cells were then plated to achieve final 60-70% confluency by the time of IF processing. Cells were washed with 1x PBS 3 times, then fixed in 4% PFA for 10 min at room temperature in the dark. PFA was washed with 1x PBS for another 3 times. Cells were then permeabilized with PBST (1x PBS with 0.25% Triton-X) for 10 min. Cells were then blocked with 5% bovine serum albumin (BSA) in 0.3% Triton-X PBST for 30 min. Cells were then processed for primary antibody staining at 4°C for overnight with a negative control without any primary antibody with indicated antibody dilution shown in the methods section. Antibody staining and washing buffer is 1% BSA in 0.3% PBST. After primary antibody staining, cells were washed for 3 times. Secondary antibody was stained at 1:400 dilution regardless of types of fluorophore conjugated secondary antibody for 1 hr at room temperature along with 1:10,000 dilution of Hoechst 33342. After secondary staining, cells were washed again for 3 times. Last, the coverslips were mounted using Vectashield Vibrance Antifade Mounting Media (H-1700-10). After overnight curing in the dark at room temperature, the cells were imaged using Zeiss LSM 980 Airyscan 2 microscope and detector equipped with a 63x oil objective with a 1.4 numerical aperture. Post-processing was done with Zen 3.4 (Blue edition).

### Proximity ligation assay

Cover glasses (#1.5H Thickness Ø12, Thor Labs) were coated with 0.1mg/ml of poly-D-Lysine (Sigma) for 1 hour at 37°C, washed once with sterile water, dry in the biosafety cabinet for 2 hours, and then complete media was added on to equilibrate for 20 min in the incubator. 25K of HT1080 and 50K of SW480 cells were then plated to achieve final 60-70% confluency by the time of imaging. PLA procedure was conducted via suggested manufacture using Duolink® In Situ Detection Reagents FarRed (Sigma-Aldrich, cat#DUO92013) with primary antibody pairs TOM20 and IP3R-I/II/III.

### Lattice Light sheet imaging

Same cell conditions were used as live cell confocal imaging. Zeiss Lattice Light Sheet 7 were used and imaged with 20x objective with an NA of 1.0. Cells were treated with Fedratinib and then monitored for 16 hr with images getting acquired every 5 min. Initial post-processing was done with Zen 3.4 (Blue edition). Images were deconvolved and then deskewed in the Zen 3.4. (Deskewing transforms the volume into traditional X, Y, Z coordinates). Then, Arivis Vision 4D (ver. 4.1.0) was used for puncta and cell segmentation. Deconvolved and deskewed datasets were analyzed with Arivis Vision 4D (ver. 4.1.0). For punctas, segmentation was done via the Blob Finder tool. The resulting segments were then filtered by size to remove false positives. For cells, maximum intensity projections of the deconvolved deskewed data were created. Hand segmentations were used to train a machine learning segmentation model. The model was then used to segment each time point in the maximum intensity projection time series. To get the number of cells per time point, the total area of the segments was divided by the average area of a cell. Puncta per cell was then further calculated.

### Transmission electron microscopy

SW480 and HT1080 cells treated with vehicle or Fedratinib for 24 hr were fixed in 3% glutaraldehyde and 3% paraformaldehyde in 0.1LM Cacodylate buffer (CB; pH 7.2) overnight at 4 °C. Cells were then washed with PBS and centrifuged at 2000 rpm for 2 mins. Pre-warmed 2% agarose solution was then carefully added to islet pellets, centrifuged and allowed to cool at 4 °C for 30 min. Samples were then subjected to osmification in 1.5% K4Fe(CN)6 + 2% OsO4 in 0.1 CB for 1 h, dehydrated by serial washes in EtOH (30%, 50%, 70%, 80%, 90%, 95% and 100%) and embedded in Spurr’s resin by polymerization at 60 °C for 24 h. Polymerized resins were then sectioned at 90 nm thickness using a Leica EM UC7 ultramicrotome and imaged at 70 kV using a JEOL 1400 TEM equipped with an AMT CMOS imaging system. Cristae classifications were done by blinding all images, and then manually annotated with the four cristae classification: laminar, non-laminar, compartmentalized, and loss-of-cristae. Mitochondrial structures (Aspect Ratio, Perimeter, Circularity) were analyzed and quantified by Fiji/ImageJ. A minimum of 150 mitochondria are analyzed from 25 independent cells.

### Correlative light and electron microscopy (CLEM)

SW480 SPLICS reporter cells were induced with doxycycline (100 ng/ml) for 24 hr, and then treated with Fedratinib for 24 hr. Cells were high pressure frozen with Leica EM High Pressure Freezer, freeze substituted using 0.1% Uranyl acetate in acetone, and finally infiltrated in Lowicryl HM20, following the protocol published in Ronchi et al., 2021. 300nm sections were cut using a Leica UC7 ultramicrotome and picked up on carbon coated 200 mesh grid**s**. Fluorescence microscopy imaging of the sections was carried out as previously described (Kukulski et al., 2012) using a wide-field Olympus IX81, equipped with an Olympus PlanApo 100X 1.4NA oil immersion objective. After that, tilt series of the areas of interest containing fluorescent signal were acquired using a TECNAI F30 (Thermo Fisher Scientific) and tomograms were reconstructed using IMOD (Kremer et al, 1996). High precision overlays between fluorescence and electron microscopy images were carried out using the plug in ec-CLEM (<u>Paul-Gilloteaux et al., 2017</u>) of the software platform Icy (<u>de Chaumont et al., 2012</u>), by clicking on corresponding features in the two imaging modalities. These articles were cited for reference^56–59^.

### Segmentation for CLEM

Automated detection of membranes in the tomogram was performed using the TomoSegMemTV (April 2020 version) software package (PMID: 24625523). The parameters used for membrane enhancement were as follows: scale_space -s 3; dtvoting -s 10 (default); surfaceness -m 0.45 -s 1.0 -p 1.0 (default); dtvoting -w -s 10 (default); surfaceness -S - m 0.3 (default); thresholding -l 0.05; global analysis −3 100 (default). The output volumes were imported into AMIRA 2022.2 (Thermo Fisher Scientific), and segmented membrane pixels were manually annotated as specific organelle membranes using the 3D magic wand tool and saved as different labels. Further, gaps in the segmentation were filled using the paint tool. The final membrane surfaces were generated using the ‘generate surface’ module and smoothened using the ‘smooth surface’ module with the following parameters: iterations 100; lambda 0.2.

### High pressure freezing, FIB-SEM, and data processing

Cells were cultured on optically flat sapphire disks (3mm diameter, 50-80um thick, Nanjing Co-Energy Optical Crystal Co., Ltd.). Cell growth media was exchanged 3x for freezing media containing Fluorobrite media (A1896702; ThermoFisher), 25% Dextran (Mr ∼40,000, 31389–100G; Sigma), and 0.8pM TetraSpeck microspheres (0.2μm diameter, T7280; Invitrogen). They were then immediately subjected to High Pressure Freezing (HPF Compact 01; Wohlwend GmbH) according to the manufacturer’s instructions. Samples were then stored in liquid N_2_. Samples were loaded onto a custom CryoSIM microscope as described in the following citation at the end of the paragraph. Epi-fluorescence montage images were taken of each coverslip to select cells for higher-resolution structured illumination microscopy (SIM). 3D SIM images were acquired (5 phases, 3 angles) over a 7-10um z-depth, and were reconstructed as described in the following citation at the end of the paragraph. Typical reconstruction parameters included 0.007 Wiener Filter, 0.7 gamma apodization, 15-pixel radii of the singularity suppression at the OTF origins. From SIM images, candidate cells for FIB-SEM were selected. Samples were subjected to a freeze-substitution/resin-embedding procedure in 2% OsO_4_, 0.1% Uranyl Acetate, and 3% water in acetone under liquid N_2_ using a Leica AFS system. Samples were infiltrated and embedded with Eponate 12 resin as described in in the following citation at the end of the paragraph. After coverslips were removed from the resin block (leaving cells embedded), repeated ultramicrotome trimming (Leica EM UC7) and micro X-ray CT imaging (Zeiss Xradia 510) was used in conjunction with cryo-fluorescence image registration to locate the cells of interest in the resin block. Samples were mounted on a copper stud, and sputter coated with 10nm gold and 100nm carbon (PECS 682; Gatan). Samples were loaded onto a custom FIB-SEM, consisting of a Zeiss Gemini 450 Field Emission SEM and a Zeiss Capella FIB column oriented at 90 degrees to the SEM beam^60^.Samples were imaged using 0.9kV landing energy and 0.25nA electron dose, and images were acquired at 200kHz readout rate (5us dwell time), and 4nm pixel size. Samples were milled using a 15nA gallium source at 30kV acceleration voltage. Actual milling time and current per slice was adjusted to maintain average 4nm milling rate, producing isotropic 4nm datasets. Images slices were registered with respect to each other using a python implementation of the SITF algorithm^61^ with RANSAC solver and a regularized affine transformation model, available at https://github.com/gleb-shtengel/FIB-SEM. Images were further de-noised using the noise 2 void 2 (N2V2) algorithm, available at https://github.com/juglab/n2v. Overall, similar workflow was performed in these cited publications^60,62,63^.

### Segmentation and analysis of FIBSEM

The ER and mitochondria were segmented using deep-learning based segmentation software Empanada (EM Panoptic Any Dimension Annotation) (PMID: 36657391). The pretrained MitoNet model was finetuned for instance-based segmentation of mitochondria. A new model was trained for ER segmentation. Semantic and instance-based segmentations were generated for ER and mitochondria, respectively. Instance-based segmentation allowed assignment of unique identifier to each mitochondrion. The mitochondrial segmentations were further improved by manually deleting the falsely segmented objects. The ER segmentations were cleaned by removing unspecific objects less than 2000 pixel2. Following this, the segmentations were further improved by manually deleting larger false positive objects. The segmentations were binned by a factor of 4 for analysis in Fiji (PMID: 22743772). Mitochondrial volumes and ERMCS distances were calculated using the 3D manager plugin. For the quantification of ER-mitochondria contact surface area, single-voxel-thick outer mitochondrial surfaces were generated. Then, the ER objects were expanded by a distance of 64 nm (4 voxels). The overlapping voxels of the expanded ER objects and mitochondria surface were marked as the contact surface. The percentage of mitochondrial contact surface area was calculated by dividing the contact surface area by the total mitochondrial surface area.

### Sample preparation for LC-MS/MS metabolomics and lipidomics analysis

Samples for metabolomics and lipidomics LC-MS/MS analyses were prepared by following the automated dual-metabolite/lipid sample preparation workflow described in the Agilent application note 5994-5065EN. Briefly, cells (ca. 1M) were collected, washed with PBS and lysed with 1:1 trifluoroethanol/water at room temperature. Lysates were transferred to microcentrifuge tubes, incubated for 10 minutes, quickly centrifuged at 250 xg for 30 seconds, and dried out by centrifugation under reduced pressure with no heat. Samples were resuspended with 1:1 trifluoroethanol/water, transferred to a 96-well plate, and processed on a Bravo Metabolomics sample preparation platform (Agilent Technologies, Inc.) with two separate VWorks protocols to sequentially and selectively isolate cell metabolites and lipids as described (5994-5065EN).

### LC-MS/MS metabolomics and lipidomics analysis

Sample were analyzed on an Agilent 1290 Infinity II Bio LC ultra-high performance liquid chromatography (UPLC) system consisting of a high-pressure binary pump, multicolumn thermostat, and a temperature controlled multisampler. The LC modules were setup with a standard configuration for omics workflows, which allowed easy acquisition method interchange for polar metabolites and lipid analyses. Samples for both targeted metabolomics and lipidomics analysis were analyzed in randomized order on an Agilent 6495C triple quadrupole mass spectrometer equipped with an Agilent Jet Stream Dual ESI ion source. For targeted metabolomics, isolated polar metabolites were analyzed with a HILIC LC-MS/MS method as described in the Agilent application note 5994-5628EN. For targeted lipidomics analysis, samples were analyzed with the reverse phase LC-MS/MS method reported in the Agilent application note 5994-3747EN. After acquisition, both metabolomics and lipidomics datasets were processed with MassHunter Quantitative analysis software and subsequently imported into Mass Profiler Professional (MPP) for chemometrics analysis. Metabolomics and lipidomics data were analyzed using MetaboAnalyst. In particular, statistical analysis [one factor] was selected. Data table with metabolite peak intensities in rows and samples in column was used. Data were filtered based on interquartile range (IQR) set filter out 25%, normalized based on protein quantitation, transformed with Log base 10, scaled by mean-centering and dividing by the standard deviation of each variable. Data were visualized with principal component analysis (PCA) and volcano plots.

### Mitochondrial lipidomics

Mitochondria were isolated with the same method detailed prior. Lipid extraction was performed as described by Bielawski^64^ with slight modifications. Samples were thawed on ice and then samples were extracted with 1.0 mL of IPA:Water:EtOAc (30:10:60, v:v:v) and internal standard mixture of EquiSPLASH™ LIPIDOMIX® Quantitative Mass Spec Internal Standard (Avanti Polar Lipids, Birmingham, AL). The extract was vortexed and sonicated 2 minutes, followed by centrifugation for 10 minutes at 8000 x g at 4 °C. The organic upper phase was transferred to a new tube. The pellet was re-extracted with an additional 0.5 mL of IPA:Water:EtOAc (30:10:60, v:v:v). The supernatants were combined and placed at -20 °C for 24 h. The supernatants were dried down using a speed vac. The dried sample was reconstituted in 150 µL of the initial condition of the mobile phase. The suspension was vortexed for 5 minutes and then centrifuged for 10 minutes at 17000 x g at 4 °C. The supernatant was transferred to an auto-sampler vial for UHPLC-MS analysis.

Lipid profiling was conducted using a Vanquish UHPLC system with an Orbitrap Fusion Lumos Tribrid™ mass spectrometer using a H-ESI™ ion source (all Thermo Fisher Scientific, Waltham, MA) with a Waters ACQUITY UPLC CSH C18 column (150 mm x 1mm, 1.7 µm particle size, Milford, MA). Solvent A was HPLC grade Water:acetonitrile (40: 60, v:v) with 0.1% formic acid and 10 mM ammonium formate. Solvent B was HPLC grade isopropanol:acetonitrile (95:5, v:v) with 0.1% formic acid and 10 mM ammonium formate. The column was maintained at 65 °C and a flow rate was set at of 110 µL/min. The gradient of the solvent B is 15 % (B) at 0 min, 30 % (B) at 2 min, 2–2.5 min 48% (B), 2.5–11 min 82 % (B), 11–11.01 min 99 % (B), 11.01–12.95 min 99 % (B), 12.95–13 min 15 % (B), and 13–15 min 15 % (B). Data acquisition was carried out in positive and negative charge modes, with the ion source spray voltage configured to 4,000 V and 3,000 V, respectively. The mass spectrometry analysis spanned a scan range of 200 to 1600 m/z for the full scan, and the MS1 resolution was established at 120K at m/z 200. The AcquireX deep scan mode was employed for the MS2 acquisition, which was performed with a stepped collision energy (SCE) of 30%, along with a 5% spread for the positive fragment ion MS/MS scan and with a SCE of 40 % along with a 20 % spread for the negative fragmental scan.

Raw data was converted into mzML format using Proteowizard mscovert software^65^. MS-DIAL software (version v5.4.241004)^66^ was used for general lipidomics data analysis including compound identification with LipidBlast^67,68^ which is default library in MS-DIAL. The data was normalized by EquiSPLASH spiked-in internal standards.

### Targeted CoQ Measurement by LC-MS/MS

Determination of the CoQ content and redox state in mammalian cell culture and isolated mitochondria (with described isolation process) was performed as previously described with modifications. In brief, frozen cell pellets or mitochondria pellets were resuspended in 200 μL of PBS and added to ice cold extraction solution (200 μL acidified methanol [0.1% HCl final], 300 μL hexane, with 0.1 μM CoQ_8_ internal standard).

Samples were vortexed and centrifuged (5 min, 17000 x g, 4°C) and the top hexane layer was transferred to a new tube. Extraction was repeated twice before the hexane layers were combined and dried under argon gas at room temperature. Extracted dried lipids were resuspended in methanol containing 2 mM ammonium formate and overlaid with argon.

LC-MS analysis was performed using a Thermo Vanquish Horizon UHPLC system coupled to a Thermo Exploris 240 Orbitrap mass spectrometer. For LC separation, a Vanquish binary pump system (Thermo Fisher Scientific) was used with a Waters Acquity CSH C18 column (100 mm × 2.1 mm, 1.7 μm particle size) held at 35 °C under 300 μL/min flow rate. Mobile phase A consisted of 5 mM ammonium acetate in acetonitrile:H_2_O (70:30, v/v) with 125 μL/L acetic acid. Mobile phase B consisted of 5 mM ammonium acetate in isopropanol:acetonitrile (90:10, v/v) with the same additive. For each sample run, mobile phase B was initially held at 2% for 2 min and then increased to 30% over 3 min. Mobile phase B was further increased to 50% over 1 min and 85% over 14 min and then raised to 99% over 1 min and held for 4 min. The column was re-equilibrated for 5 min at 2% B before the next injection. Five microliters of the sample were injected by a Vanquish Split Sampler HT autosampler (Thermo Fisher Scientific), while the autosampler temperature was kept at 4 °C. The samples were ionized by a heated ESI source kept at a vaporizer temperature of 350 °C. Sheath gas was set to 50 units, auxiliary gas to 8 units, sweep gas to 1 unit, and the spray voltage was set to 3500 V for positive mode and 2500 V for negative mode. The inlet ion transfer tube temperature was kept at 325 °C with 70% RF lens. For targeted analysis, the MS was operated in parallel reaction monitoring mode with polarity switching acquiring scheduled, targeted scans to oxidized CoQ_10_ H+ adduct (m/z 863.6912), oxidized CoQ_10_ NH+ adduct (m/z 880.7177), reduced CoQ_10_H_2_ H+ adduct (m/z 865.7068), reduced CoQ_10_H_2_ NH+ adduct (m/z 882.7334), CoQ_8_ H+ adduct (m/z 727.566) and CoQ intermediates: DMQ_10_ H+ adduct (m/z 833.6806), and PPHB_10_ H-adduct (m/z 817.6504). MS acquisition parameters include resolution of 15,000, HCD collision energy (30% for positive mode and stepped 20%, 40%, 60% for negative mode), and 3s dynamic exclusion. Automatic gain control targets were set to standard mode. The resulting CoQ intermediate data were processed using TraceFinder 5.1 (Thermo Fisher Scientific). Raw intensity values were normalized to the CoQ_8_ internal standard and protein content as determined by BCA^69^.

### FACS sorted mitochondrial protein proteomics

All samples were lysed in lysis buffer (8M Urea, 4-(2-Hydroxyethyl)-1-piperazinepropanesulfonic acid (EPPS)) with sonication. Samples were precipitated with SP3 beads and labeled with iodoacetamide, washed and resuspended in 100mM EPPS buffer, pH 8.5 and digested at 37C with trypsin/LysC overnight. The samples were labeled with TMT Pro and quenched with hydroxylamine. All the samples were combined and desalted using a 50 mg Sep-Pak cartridge, followed by drying in a speedvac. Samples were desalted via StageTip and dried with speedvac. Samples were resuspended in 5% formic acid, and 5% acetonitrile for LC-MS/MS analysis.

Mass spectrometry data were collected using a Orbitrap Astral mass spectrometer (Thermo Fisher Scientific, San Jose, CA) coupled with Neo Vanquish liquid chromatograph. Peptides were separated on a 110 cm uPAC C18 column (Thermo Fisher Scientific). For each analysis, we loaded ∼0.5 μg onto the column. Peptides were separated using a 150 min gradient of 5 to 29% acetonitrile in 0.125% formic acid with a flow rate of 300 μL/min.

The scan sequence began with an Orbitrap MS^1^ spectrum with the following parameters: resolution 120,000, scan range 350−1350 Th, automatic gain control (AGC) target 200%, maximum injection time 50ms, RF lens setting 50%, and centroid spectrum data type. FAIMS was enabled with compensation voltages (CVs): -35V, -45V, -55V, -60V, and -70V using TopSpeed setting of 1 sec for each CV. We selected the top twenty precursors for MS^2^ analysis which consisted of HCD high-energy collision dissociation with the following parameters: Astral data acquisition (TMT off), AGC 200%, maximum injection time 15ms, isolation window 1.2 Th, normalized collision energy (NCE) 36, and centroid spectrum data type. In addition, unassigned and singly charged species were excluded from MS^2^ analysis and dynamic exclusion was set to 90 s.

### Whole cell proteomics

All samples were labeled with iodoacetamide and resuspended in 100 mM of 4-(2-Hydroxyethyl)-1-piperazinepropanesulfonic acid (EPPS) buffer, pH 8.5 and digested at 37C with trypsin/LysC overnight. The samples were labeled with TMT Pro and quenched with hydroxylamine. All the samples were combined and desalted using a 100mg Sep-Pak cartridge, followed by drying in a speedvac. Samples were fractionated with basic pH reversed phase (BPRP) high-performance liquid chromatography (HPLC) as described previously^70^. Samples were desalted via StageTip and dried with a speedvac. Samples were resuspended in 5% formic acid, and 5% acetonitrile for LC-MS/MS analysis. The mass spectrometry proteomics data have been deposited to the ProteomeXchange Consortium via the PRIDE partner repository with the dataset identifier PXD056917.

### Liquid chromatography and tandem mass spectrometry

Mass spectrometric data were collected on an Exploris480 mass spectrometer coupled to a Proxeon NanoLC-1200 UHPLC. The 100 µm capillary column was packed with 35 cm of Accucore 150 resin (2.6 μm, 150Å; ThermoFisher Scientific) at a flow rate of 450 nL/min for 90min. The scan sequence began with an MS1 spectrum (Orbitrap analysis, resolution 60,000, 350-1350 Th, automatic gain control (AGC) target is set to “standard”, maximum injection time set to “auto”). Data were acquired for 150 minutes per analysis. The hrMS2 stage consisted of fragmentation by higher energy collisional dissociation (HCD, normalized collision energy 32%) and analysis using the Orbitrap (AGC 300%, maximum injection time 96 ms, isolation window 0.7 Th, resolution 30,000 with TurboTMT activated). Data were acquired using the FAIMSpro interface the dispersion voltage (DV) set to 5,000V with compensation voltages (CVs) set at -40V, -60V, and -80V. The TopSpeed parameter was set at 1 sec per CV.

### Proteomics data analysis

Raw files were searched using the Comet algorithm with a custom database search engine reported previously^71^. Database searching included human (Homo Sapiens) entries from UniProt (http://www.uniprot.org, downloaded 2021) with the reversed sequences, and common contaminants (*i.e.,* keratins, trypsin). Peptides were searched using the following parameters: 50 ppm precursor mass tolerance; up to 2 missed cleavages; variable modifications: oxidation of methionine (+15.9949); static modifications: TMTpro (+304.2071) on lysine and peptide N terminus, carboxyamidomethylation (+57.0215) on cysteines. The protein-level FDR was determined using the ModScore algorithm where a score of 13 corresponds to 95% confidence in correct localization. TMT reporter ions were used for quantification of peptide abundance. Isotopic impurities were corrected according to the manufacturer’s specifications, and signal-to-noise (S/N) was calculated. For Astral MS data, samples were processed by removing peptides with summed S/N lower than 1000 across all channels or isolation specificity lower than 0.7. Spectra with resolution <45,000 were discarded. For samples acquired with Exploris 480, peptides with summed S/N lower than 100 across all channels or isolation specificity lower than 0.7 were excluded. The high confidence peptides were then used to quantify protein abundance by summing up S/N values for all peptides assigned to the same protein, and only proteins in the linear quantification range of the instrument were included in the analysis. The normalization for protein quantification was then performed by adjusting protein loadings of total sum of S/N values to that of every TMT channel.

### Chromatin immunoprecipitation (ChIP)

Chromatin immunoprecipitation (ChIP) experiments were performed using the Ideal ChIP-seq kit for transcription factors (Diagenode) as per the manufacturer’s protocol. 4 million SW480 and HT1080 cells treated with vehicle (DMSO), thapsigargin or fedratinib were used per ChIP reaction with 8ug of the BRD4 antibody (Diagenode, Cat# C15410337). Briefly, cells were crosslinked for 10 minutes in a 1% formaldehyde solution, followed by termination with 1/10th the volume of 1.25M glycine for 5 minutes at room temperature. Following this, the cells were lysed and sonicated (Bioruptor Pico, Diagenode) to obtain desired chromatin fragments of about 200bp. Sheared chromatin was then incubated with the BRD4 antibody overnight at 4 °C. Next day, ChIP-DNA was de-crosslinked and purified using the Diagenode iPure Kit V3, following the manufacturer’s protocol. Utilizing the manufacturer’s instructions (Illumina), purified DNA was prepared for sequencing. About 1-10ng of ChIP DNA were converted to blunt-ended fragments using T4 DNA polymerase, *Escherichia coli* DNA polymerase I large fragment (Klenow polymerase), and T4 polynucleotide kinase (New England BioLabs (NEB)). Klenow fragment (3′ to 5′ exo minus; NEB) was used to add a single adenine base to fragment ends, followed by ligation of Illumina adaptors (Quick ligase, NEB). PCR enriched the adaptor-ligated DNA fragments using the Illumina Barcode primers and Phusion DNA polymerase (NEB). PCR products were size-selected using 3% NuSieve agarose gels (Lonza) followed by gel extraction using QIAEX II reagents (Qiagen). Quantified libraries quality-checked using the Bioanalyzer 2100 (Agilent) and sequenced on the Illumina HiSeq 2500 Sequencer (125-nucleotide read length). Reads were first processed using Trimmomatic version 0.39 (settings TruSeq3-PE-2.fa:2:30:10, minlen 50) followed by alignment with bwa (“bwa mem,” options -5SP -T0, version 0.7.17-r1198-dirty) to hg38 (GRCh38) reference^72,73^. After alignment the reads were filtered using Markduplicates from Picard and then by quality score of >20 via samtools. MACS2 was used to call peaks, filtered using bedtools, and converted to bigwigs with UCSC wigtoBigwig^74,75^. Cistrome overlap analysis was performed in R (v3.6.0) using ChipSeekAnno (v3.0.0) and ChipSeeker (v1.29.1)^76,77^. Enrichment heatmaps were generated using Deeptools.

### RNA Extraction, Reverse Transcription, Real-time PCR and Bromouridine nascent mRNA-Sequencing

RNA was extracted from SW480 and HT1080 cells using the QIAGEN RNeasy mini kit according to the manufacturer’s instructions. 1 ug of RNA was reverse transcribed using the Lunascript Supermix (NEB) according to the manufacturer’s instructions. Quantitative real-time PCR (qPCR) from cDNA was performed using SYBR green master mix and beta actin (ACTB) was used as a control. qPCR from BRD4 ChIP-DNA was performed using 1% of input genomic DNA with two sets of primers targeting MYC PVT enhancer locus, and a control primer set targeting an upstream region that does not have BRD4 occupancy. The primer sequences are listed in Supplementary Table 5. BrU-Seq method is described in these papers^78,79^.

### Reproducibility

Each cell line experiment was performed in technical replicates for each condition and repeated at least three times with biological triplicates to ensure reproducibility. Figures show a representative biological replicate unless otherwise indicated. Blinding was performed whenever appropriate. Sample description and identification was unavailable to the core personnel during data collection and analysis. Statistical details of all experiments can be found in the figure legends. The sample numbers are mentioned in each figure legend and denote biological replicates. Statistical details are reported in figure legends. Results are expressed as the mean plus or minus the standard error of the mean for all figures unless otherwise noted. Significance between 2 groups was tested using a 2 tailed unpaired t test. Significance among multiple groups was tested using a one-way ANOVA. GraphPad Prism 7.0 was used for the statistical analysis. Statistical significance is described in the figure legends as: ∗ p < 0.05, ∗∗ p < 0.01, ∗∗∗ p < 0.005, ∗∗∗∗ p < 0.001.

## Data availability

Images and details of the image-based drug repurposing screen were deposited to Zenodo and are publicly available in the repository as part of this record: DOI: <u>10.5281/zenodo.13963526</u>. Datasets generated are also available from the corresponding author upon reasonable request. BrU-seq data has been deposited to NCBI GEO under the accession number: GSE278077. ChIP-seq data has been deposited to NCBI GEO under the accession number: GSE280900. The mass spectrometry proteomics data have been deposited to the ProteomeXchange Consortium via the PRIDE partner repository with the dataset identifier PXD056917. Due to the size of the FIBSEM data, it will be available upon request.

## Figure Illustrations

Figure were created using Adobe Illustrator and Biorender.com.

## Drug treatment

Small molecule inhibitors used in this studied are listed in the following table. Concentration of each compound is listed in figure legend other than Fedratinib. Fedratinib was used at 1 μM for all studies unless indicated differently.

| Compound | Company | Catalog number |
| --- | --- | --- |
| Fedratinib | MedChem Express | HY-10309 |
| Thapsigargin | Cayman Chemicals | 10522 |
| Ruxolitinib | Cayman Chemicals | 11609 |
| dBET6 | Cayman Chemicals | 35385 |
| JQ1 | Cayman Chemicals | 11187 |
| Actinomycin D | Cayman Chemicals | 11421 |
| AS-2863619 | Cayman Chemicals | 30976 |
| NVP2 | Cayman Chemicals | 34725 |
| Piericidin A | Cayman Chemicals | 15379 |
| TTFA | Cayman Chemicals | 15517 |
| Antimycin A | Cayman Chemicals | 34799 |
| Myxothiazol | Sigma Aldrich | T5580 |
| Oligomycin | Cayman Chemicals | 11341 |
| CCCP | Cayman Chemicals | 25458 |
| FG4592 | Cayman Chemicals | 15294 |
| MitoPQ | Cayman Chemicals | 18808 |
| S3QEL2 | Cayman Chemicals | 18556 |
| Z-VAD-FMK | Cayman Chemicals | 14463 |
| PKR-IN-C16 | MedChem Express | HY-13977A |
| Gamitrinib TPP hexafluorophosphate (GTPP) | MedChem Express | HY-102007A |

## Acknowledgments

We thank all members of the Shah and Lyssiotis labs for their advice and suggestions. In collaboration with this research, we acknowledge support from the University of Michigan Biomedical Research Core Facilities (Flow Cytometry core, Advanced Genomics core, Microscopy core), and Bru-seq Lab. FIB-SEM imaging were performed in collaboration with the Advanced Imaging Center (AIC) at Janelia Research Campus, which is supported by the Howard Hughes Medical Institute. The funders (Agilent Technologies) for the whole cell metabolomics and lipidomics had no role in study design, data collection and analysis, or the content and publication of this manuscript. We acknowledge the European Molecular Biology Laboratory (EMBL) Electron Microscopy Core Facilities’ assistance, data acquisition, and analysis for CLEM expertise. B.C was supported by NCI F99CA284256-01. D.C.S. was supported by NIGMS 5T32GM145304-03. Y.M.S. was supported by NCI R01CA148828, R01CA245546, and NIDDK R01DK095201. C.A.L. was supported by NCI R37CA237421, R01CA248160, and R01CA244931. S.M. was supported by NIGMS DP2GM150019. D.J.P. was supported by NIDDK R01DK098672 and NIGMS R35GM131795. N.J.R. was supported by NIGMS T32GM145470 and T32GM150581. R.B. was supported by NIGMS R35GM130183. D.A.H. was supported by NIGMS F32GM140694. A.D.P. was supported by NIH S10OD021750, USDA National Institute of Food and Federal Appropriations Project PEN04917, Accession 7006712 and Pennsylvania Department of Health using Tobacco CURE funds. T.C was supported by Italian Ministry of University and Research PRIN2017, University of Padova STARS Consolidator Grant 2019 and Progetto di Ateneo 2023 no. CALI_BIRD23_01, and PNRR – CN3 National Center for Gene Therapy and Drugs based on RNA Technology n. CN00000041 (2022-26).

## Author contributions

B.C., C.A.L., and Y.M.S. designed the study and wrote the manuscript. B.C. designed experiments and collected data for the bulk of the experimental studies. B.C., D.C.S, P.J., T.M.L., B.S.H., N.J.R., N.S, M.S., J.A.P, M.C., M. P., S.K., I.K., S.E., P.R, P.M., D.A.H., J.L. R.M.G., and carried out experimental aspects of the project, and contributed to data analysis. B.C., S.M., D.J.P., N.J.R., R.B., D.A.H., M. L., A.P., J.D.M., T.C., C.A.L., and Y.M.S. provided resources, funding, and conceptual input for experiments and supervised the research. All authors are involved in the throughout the research process, agreed amongst authors regarding roles and responsibilities, and contributed to review, editing, and approval of the manuscript.

## Materials & Correspondence

Supplementary Information is available for this paper. All data generated or analyzed during this study are included in this published article and its supplementary information files. Correspondence and requests for materials should be addressed to Yatrik M. Shah or Costas A. Lyssiotis.

---

**Extended Data Figure 1:**
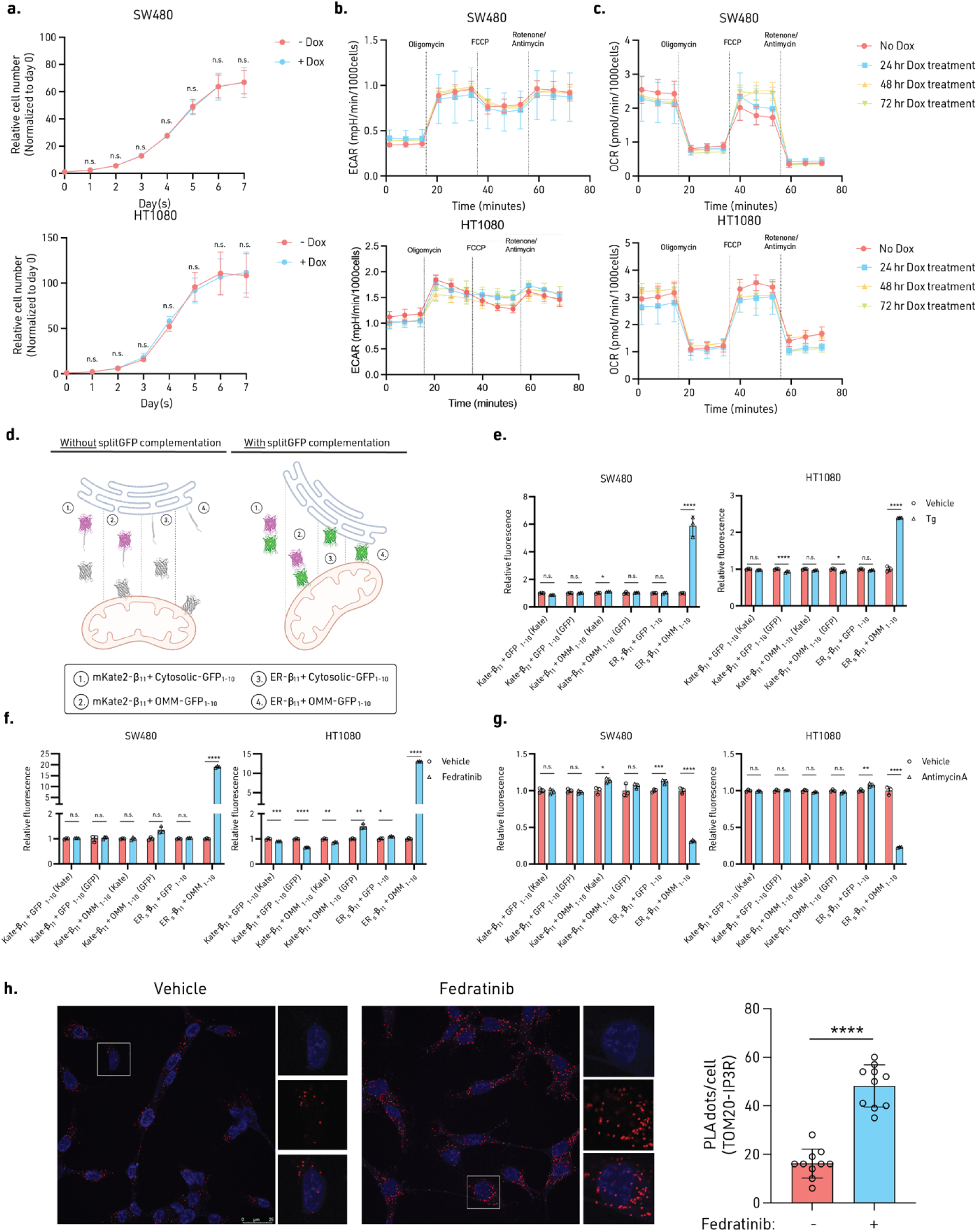
Control experiments for validating SPLICS reporter and orthogonal validation. **a**, Cell proliferation data of SPLICS reporter cells treated with doxycycline to induce reporter expression. **b**, ECAR measurement in SPLICS cells treated with 24, 48, and 72 hr of doxycycline to induce reporter expression. **c**, OCR measurement in SPLICS cells treated with 24, 48, and 72 hr of doxycycline to induce reporter expression. **d**, Schematic of control pairs for SPLICS. 1. mKate2 tagged beta_11_ + cytosolic GFP_1-10_ 2. mKate2 tagged beta_11_ + OMM-GFP_1-10_ 3. ER-beta_11_ + cytosolic GFP_1-10_ 4. ER-beta_11_ + OMM-GFP_1-10_. **e**, Relative fluorescence of SPLICS and control pairs upon thapsigargin treatment. **f**, Relative fluorescence of SPLICS and control pairs upon fedratinib treatment. **g**, Relative fluorescence of SPLICS and control pairs upon Antimycin A treatment. **h**, Representative images of PLA experiment in HT1080 cells treated with vehicle or fedratinib for 24 hr. Anti-TOM20 and Anti-IP3R were used as primary antibodies in the assay (Left). Quantification of average number of PLA foci per cell (Right). 10 cells were quantified per condition per independent experiment. Experiments (**a**-**c**, **e**) were performed 3 times, and experiment (**g**) was performed twice. n = 3. n.s. = not significant, *P ≤ 0.05, **P ≤ 0.01, ***P ≤ 0.005, ****P ≤ 0.001. Data are mean ± SD. Statistical source data are provided in Source Data Extended Data Fig. 2.

**Extended Data Figure 2:**
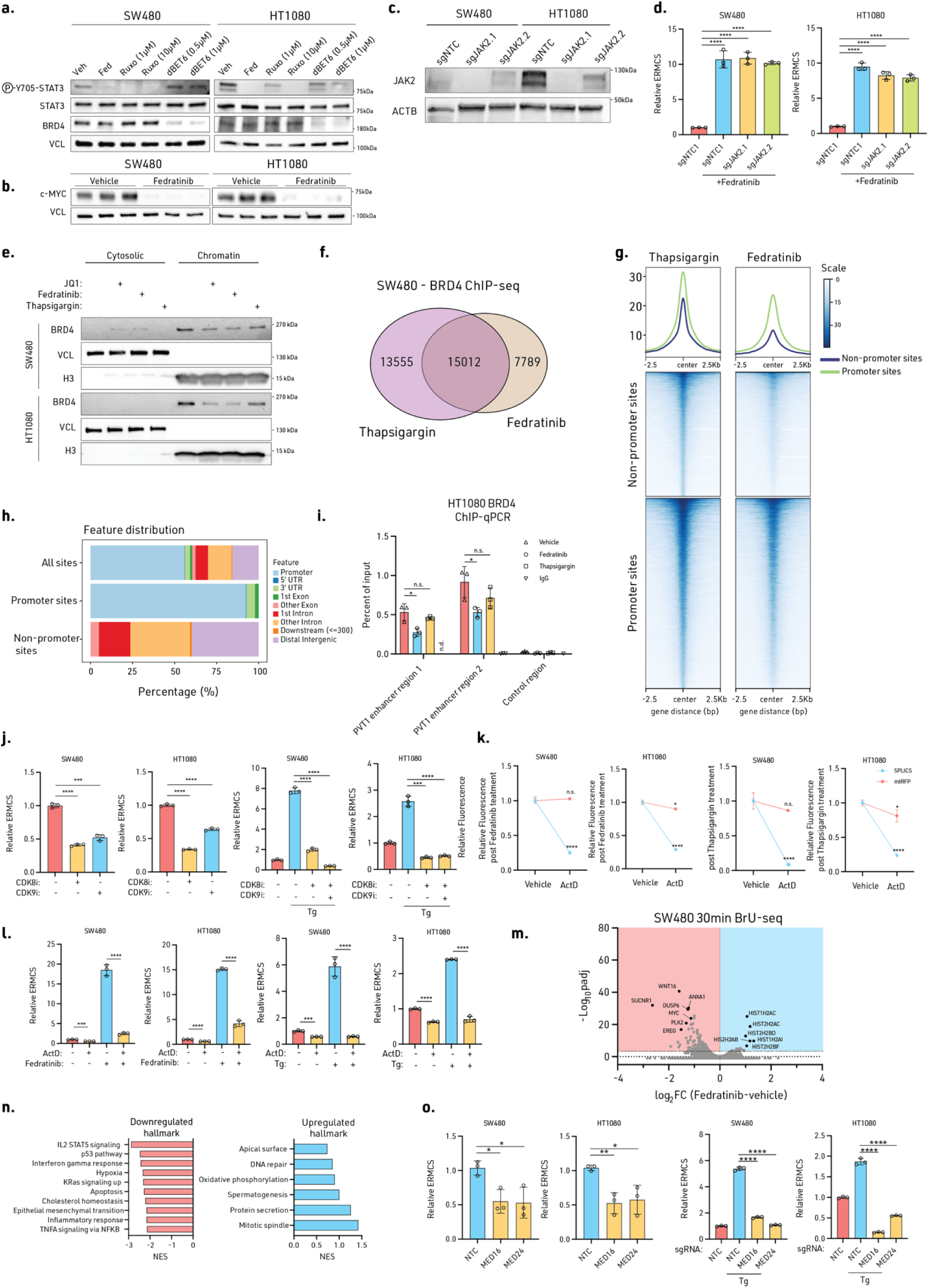
Transcriptional and epigenetic regulation of ERMCS. **a**, **b** Immunoblot from cells treated with vehicle, fedratinib (Fed), ruxolitinib (Ruxo), or dBET6 for 24 hr. **c**, Immunoblot of JAK2 KD with CRISPRi in SW480 and HT1080 cells. **d**, Relative ERMCS of control and JAK2 CRISPRi-KD cells. **e**, Chromatin fractionation assay of cells treated with JQ1 (1 μM), fedratinib, and thapsigargin (50 nM) for 24 hr. **f**, Venn diagram showing overlaps of BRD4 ChIP-seq in thapsigargin (50nM) vs fedratinib-treated SW480 cells. **g**, ChIP-seq read density heatmaps of BRD4 at promoter and non-promoter sites in thapsigargin (50 nM) vs fedratinib-treated SW480 cells. **h**, Genomic locations of BRD4 bound sites defined from the overlap analysis in panel g. **i**, BRD4 ChIP-qPCR analysis of BRD4-bound PVT1 enhancer regions in HT1080 cells treated with fedratinib or thapsigargin (50 nM) for 4 hr. **j**, Relative ERMCS in cells treated with inhibitors of transcription kinase CDK8 (AS-2863619, 1 μM) or CDK9 (NVP2, 1 μM) for 6 hr, and then treated with vehicle or thapsigargin (50 nM) for 24 hr. **k**, Actinomycin D (10 μg/ml) pulse chase of mitoTagRFP and SPLICS fluorescence after vehicle, fedratinib, or thapsigargin (50nM) treatment. **l**, Relative ERMCS fluorescence from cells treated with actimonycin D (10 μg/ml) for 24 hr, and then treated with vehicle, fedratinib, or thapsigargin (50 nM) for 24 hr. **m**, Volcano plot of BrU-seq analysis after SW480 cells were treated with fedratinib for 30 min. **n**, GSEA of BrU-seq analysis after SW480 cells were treated with fedratinib for 30 min. Normalized enrichment score (NES) of top pathways altered. **o**, Relative ERMCS and gene expression level of MED16 and MED24 mRNA transcripts upon CRISPRi mediated knockdown of cells treated with vehicle or thapsigargin (50 nM) for 24 hr. Experiments (**a**-**f**, **j-l, o**) were performed three times, and experiment (**g-i**, **m**, **n**) was performed once with biological duplicates. Significance was calculated with an unpaired two-tailed t-test. n.s. = not significant, *P ≤ 0.05, **P ≤ 0.01, ***P ≤ 0.005, ****P ≤ 0.001. Data are mean ± SD. Statistical source data are provided in Source Data Extended Data Fig. 2. Immunoblot source data are provided in Supplemental Figure 2.

**Extended Data Figure 3.**
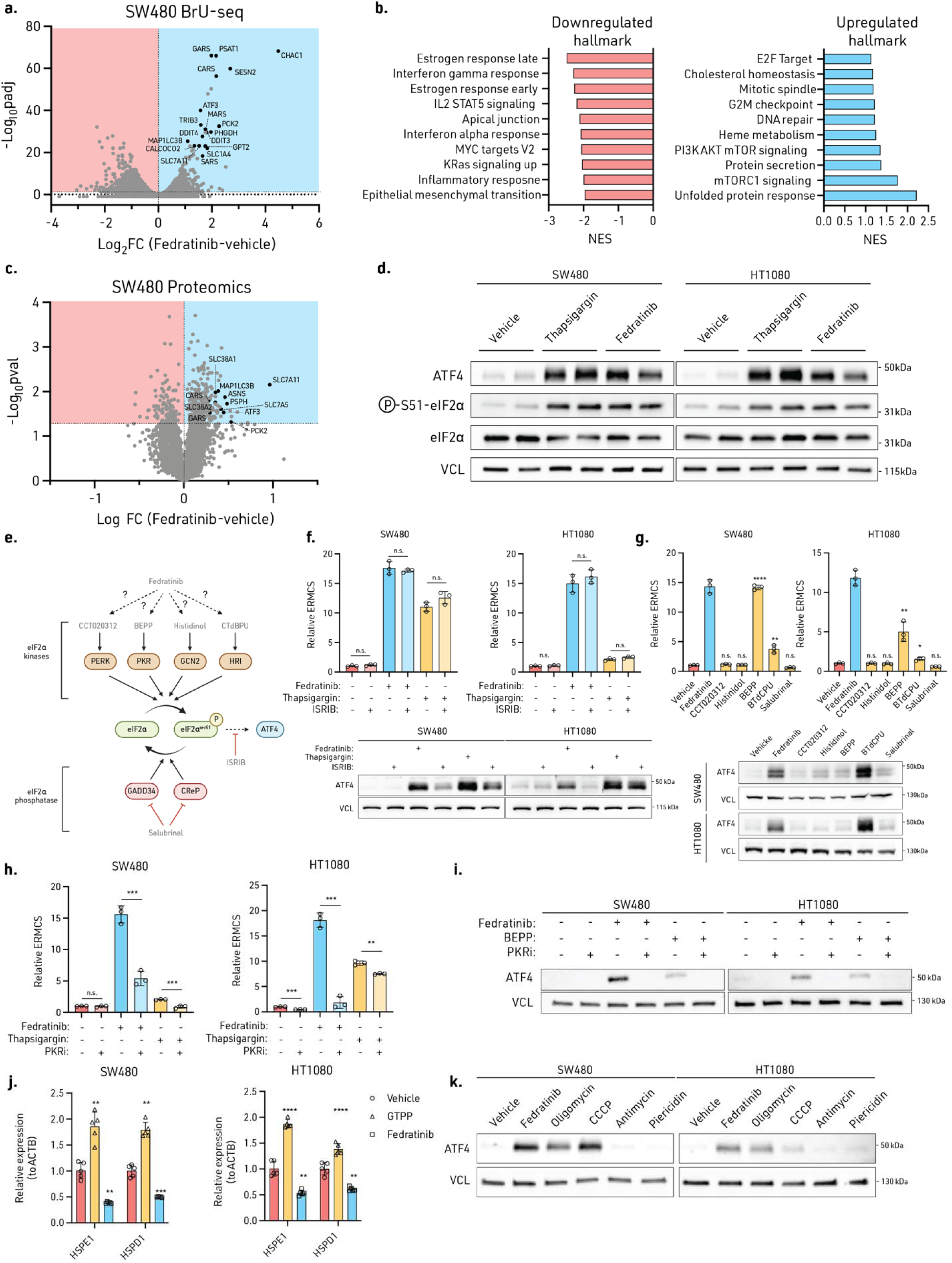
Fedratinib induces ISR and PKR supports ERMCS induction. **a**, BrU-Seq volcano plot of SW480 cells treated with fedratinib for 2 hr with ISR target genes highlighted. **b**, GSEA analysis of BrU-seq analysis after SW480 cells were treated with fedratinib for 2 hr with ISR target genes highlighted. Normalized enrichment score (NES) of top pathways altered. **c**, Proteomic volcano plot of SW480 cells treated with fedratinib for 24 hr with ISR target genes highlighted. **d**, Immunoblot of ISR targets from cells treated with fedratinib and thapsigargin (50 nM) for 24 hr. **e**, Infographic of ISR delineating the ISR kinases and phosphatases that mediate ATF4-dependent response. **f**, Relative ERMCS and ATF4 immunoblot of cells pretreated with ISRIB (1 μM) for 6 hr and then treated with fedratinib or thapsigargin (50 nM) for another 18 hr. **g**, Relative ERMCS and ATF4 immunoblot of cells treated with fedratinib and ISR inducers (CT020312, Histidinol, BEPP, BTdCPU, Salubrinal. **h**, Relative ERMCS induction and **i**, immunoblot in cells pretreated with PKR inhibitor (PKR-IN-C16, 1 μM) and caspase inhibitor Z-VAD-FMK (10 μM) for 6 hr then treated with fedratinib or thapsigargin (50 nM) for another 18 hr. **j**, Relative expression of UPRmito genes HSPD1 and HSPE1 upon fedratinib or GTPP (10 μM). **k**, Comparison of ATF4 induction via immunoblot of fedratinib to mitochondrial ETC inhibitors: Oligomycin (5 μM), Antimycin A (50 nM), Piericidin (20 nM) and uncoupler CCCP (1 μM). Experiments (**b**, **d**, **f**-**k**) were performed three times, experiments (**a**) was performed once with biological duplicates, and experiment (**c**) was performed once with four biological replicates. Significance was calculated with an unpaired two-tailed t-test. n.s. = not significant, *P ≤ 0.05, **P ≤ 0.01, ***P ≤ 0.005, ****P ≤ 0.001. Data are mean ± SD. Statistical source data are provided in Source Data Extended Data Fig. 3. Immunoblot source data are provided in Supplemental Figure 3.

**Extended Data Figure 4:**
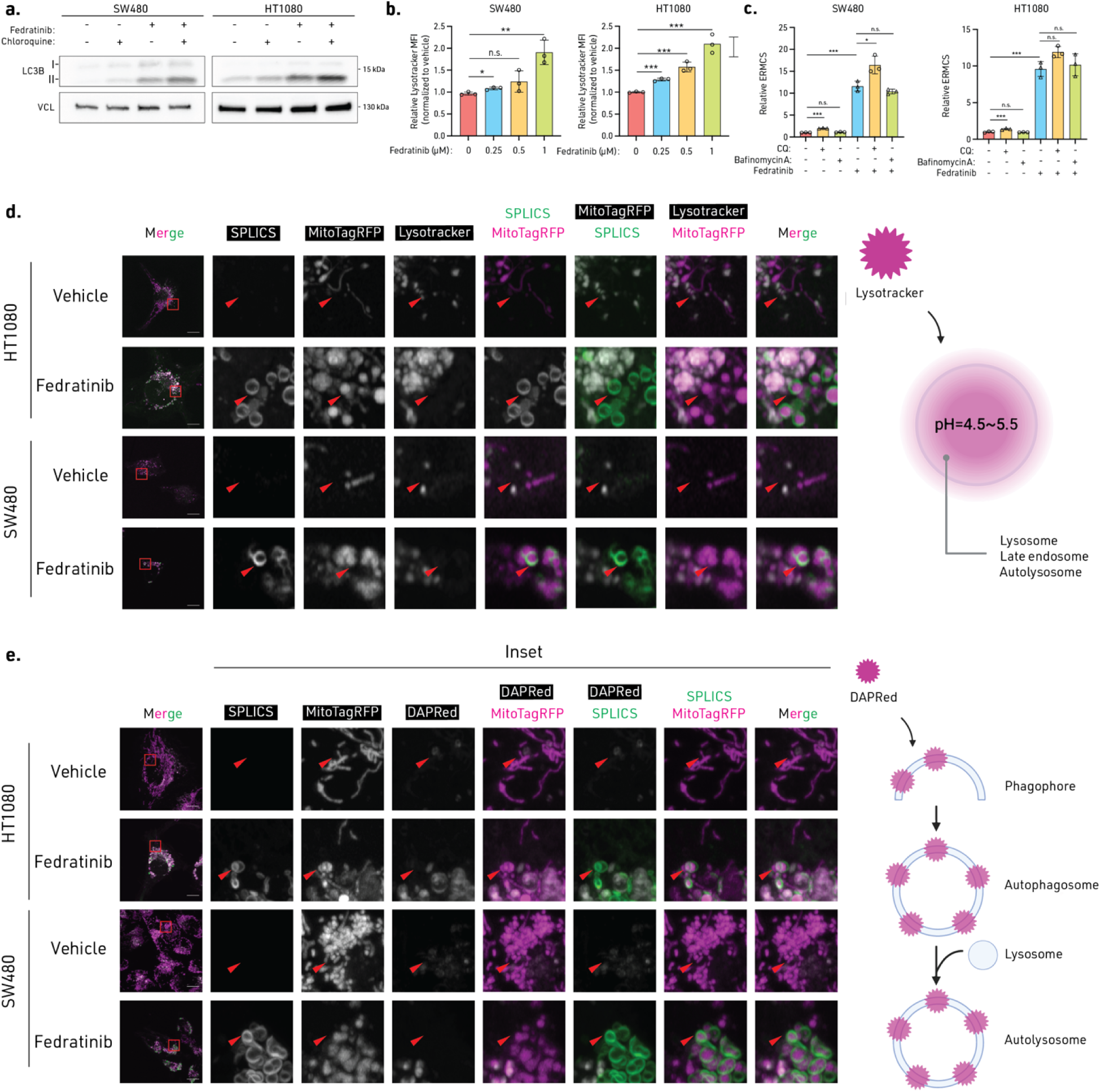
Fedratinib-dependent ERMCS formation is independent of its ability to induce autophagy. **a**, Immunoblot of delipidated (I) and lipidated (II) LC3B from fedratinib, chloroquine (CQ) and fedratinib/CQ double treated cells. **b**, Relative MFI of Lysotracker deep red in cells treated with dose dependent increase of fedratinib. **c**, Relative ERMCS from cells pretreated with CQ or bafilomycin A for 6 hr then 18 hr of fedratinib. **d**, Fluorescent images of cells treated with fedratinib with SPLICS, mitoTagRFP, and Lysotracker deep red. **e**, Fluorescent images of cells treate with fedratinib with SPLICS, mitoTagRFP, and DAPRed (autophagosome). Experiments (**a**-**c**) were performed three times, and experiment (**d**, **e**) were performed twice. Significance was calculated with an unpaired two-tailed t-test. n.s. = not significant, *P ≤ 0.05, **P ≤ 0.01, ***P ≤ 0.005, ****P ≤ 0.001. Data are mean ± SD. Statistical source data are provided in Source Data Extended Data Fig. 4. Immunoblot source data are provided in Supplemental Figure 4.

**Extended Data Figure 5:**
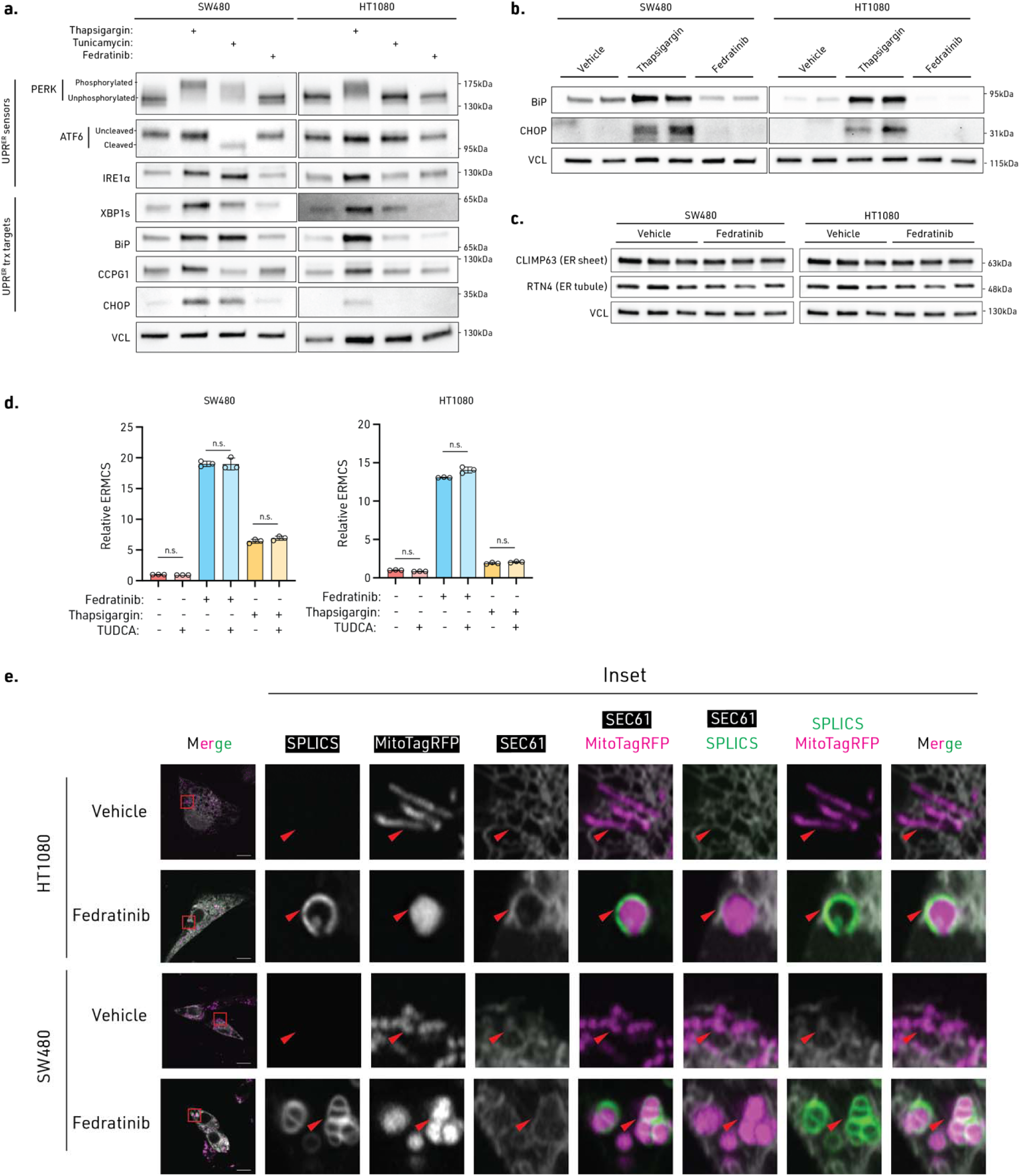
Fedratinib does not induce ER structural changes, UPR, and ER stress. **a**, **b**, **c**, Immunoblot of cells treated with fedratinib or thapsigargin (50 nM) for 24 hr for (**a**) UPR^ER^ sensors and transcription targets, (**b**) BiP and CHOP, (**c**) ER sheet and tube shaping proteins. **d**, Relative ERMCS in cells pre-treated with ER stress chaperone tauroursodeoxycholic acid (TUDCA) (400 μM) for 6 hr, and then treated with fedratinib or thapsigargin (50 nM) for another 18 hr. **e**, Microscopy images of SPLICS/mtRFP reporter cells transfected with ER marker SEC61 treated with fedratinib or vehicle control for 24 hr. Experiments (**a**-**d**) were performed three times, and experiment (**e**) was performed twice. Significance was calculated with an unpaired two-tailed t-test. n.s. = not significant. Statistical source data are provided in Source Data Extended Data Fig. 5. Immunoblot source data are provided in Supplemental Figure 5.

**Extended Data Figure 6:**
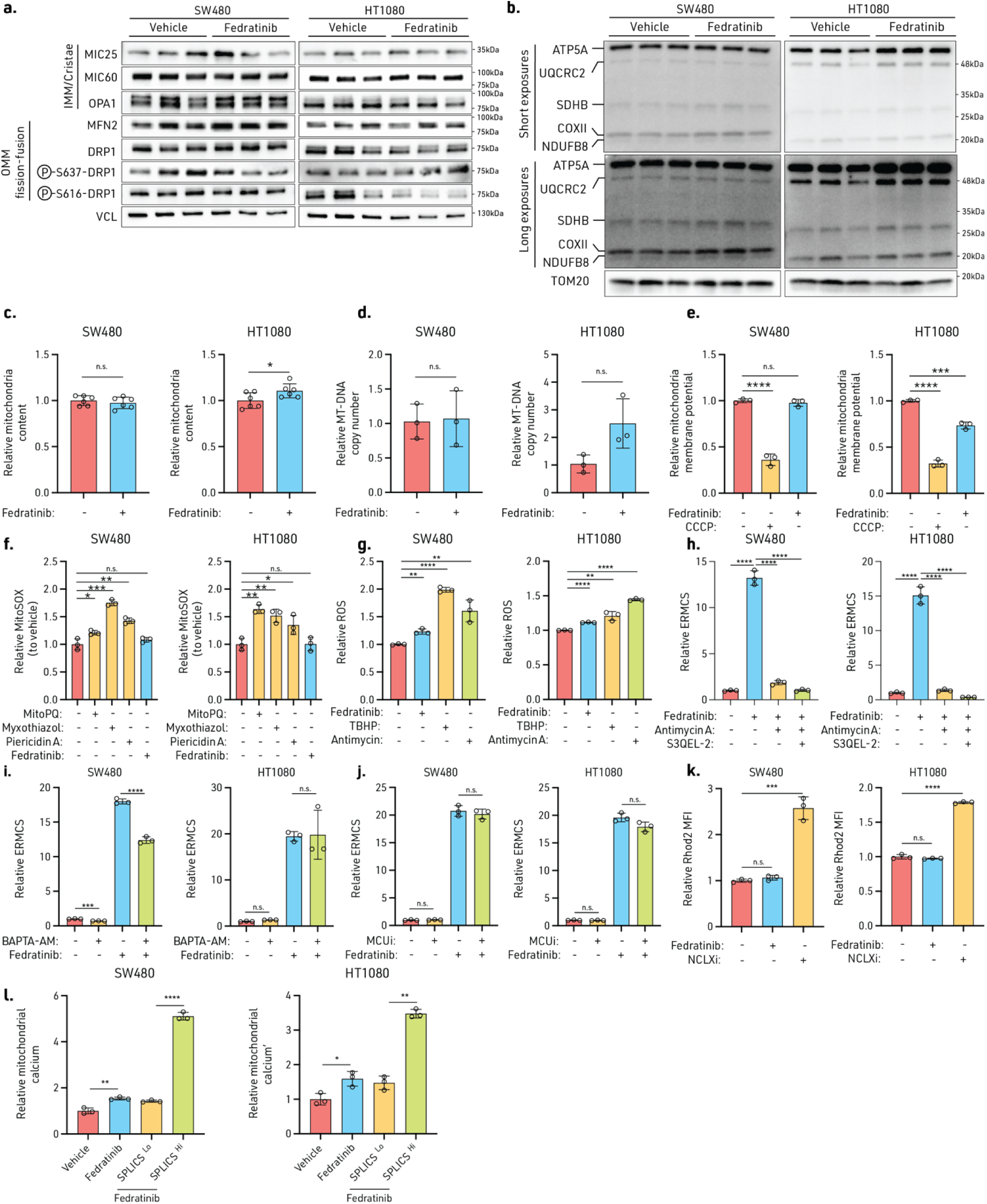
Effects of fedratfnib on various mitochondrial function. **a**, Immunoblot analyses of mitochondrial fusion (MFN2), fission (DRP1, DRP1^pS637^, DRP1^pS616^), mitochondria contact site and cristae organization system (MICOS) components (MIC25/CHCHD6, MIC60/IMMT) from cells treated with vehicle or fedratinib for 24 hr. **b**, Immnuoblot analyses of mitochondria complex I (NDUFB8), complex II (SDHB), complex III (UQCRC2), complex IV (COXII), and complex V (ATP5A) from cells treated with vehicle and fedratinib for 24 hr. **c**, Relative mitochondrial content, **d**, mitochondrial DNA (MT-DNA) copy number, **e**, mitochondrial membrane potential from cells treated with vehicle and fedratinib for 24 hr (Uncoupler CCCP (1 μM) was used as positive control). **f**, Mitochondrial ROS (mtROS) was assessed using a ratio of MitoSOX Deep Red-to-MitoTracker Green. mito-paraquat (mitoPQ, 10 μM), myxothiazol (25 nM), and piericidin A (20 nM) were used to induce mtROS. **g**, General ROS MFI was measured with CellROX Deep Red with positive control tert butyl hydrogen peroxide (TBHP, 200 μM), positive control Antimycin A (50 nM), and fedratinib for 24hr. **h**, Relative ERMCS from cells pretreated with ETC CIII ROS quencher S3QEL2 (10 μM) for 3 hr then treated with Antimycin A (50 nM) for 3 hr, and then treated with fedratinib for 18 hr. **i**, Relative ERMCS from cells treated with calcium chelation (BAPTA-AM; 5 μM) and **j**, MCU inhibition (Ru360; 10 μM) for 24 hr, and then treated with vehicle or fedratinib for 24 hr following. **k**, Mitochondria calcium was monitored with Rhod2 staining from cells treated with vehicle, positive control NCLX inhibitor, CGP-37157 (50 μM), or fedratinib for 24 hr. Experiments (**a**-**l**) were performed three times. Significance was calculated with an unpaired two-tailed t-test. n.s. = not significant, *P ≤ 0.05, **P ≤ 0.01, ***P ≤ 0.005, ****P ≤ 0.0001. Data are mean ± SD. Statistical source data are provided in Source Data Extended Data Fig. 6. Immunoblot source data are provided in Supplemental Figure 6.

**Extended Data Figure 7:**
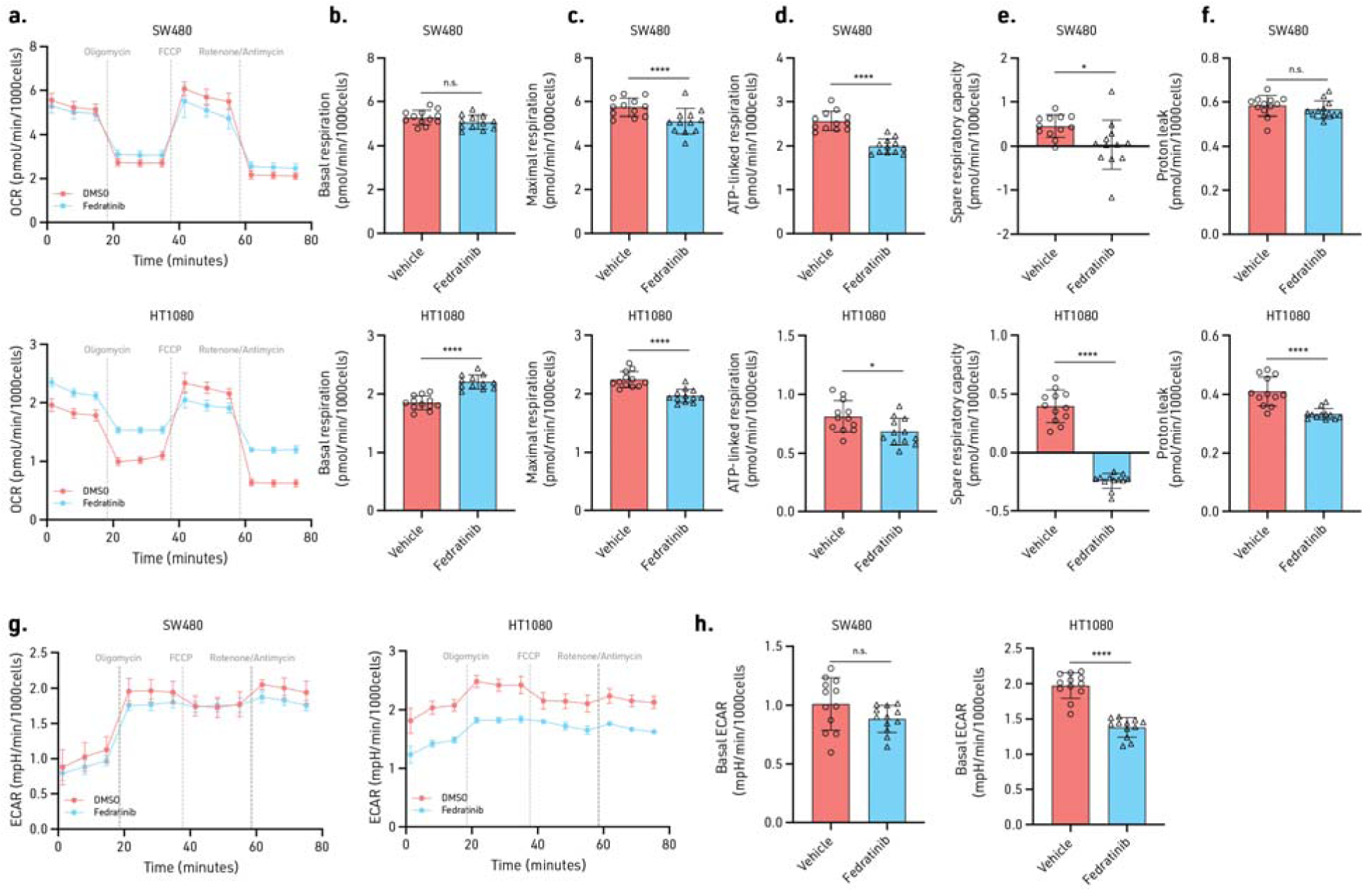
The effects of fedratinib on mitochondrial bioenergetics. **a**, Oxygen consumption (OCR), **b**, basal respiration, **c**, maximal respiration, **d**, ATP-linked respiration, **e**, spare respiratory capacity, **f**, proton leak, **g**, extracellular acidification rate (ECAR) and, **h**, basal ECAR of fedratinib or vehicle treated cells undergoing Seahorse MitoStress test. Experiments (**a**-**g**) were performed three times. Significance was calculated with an unpaired two-tailed t-test. n.s. = not significant, *P ≤ 0.05, **P ≤ 0.01, ***P ≤ 0.005, ****P ≤ 0.001. Data are mean ± SD. Statistical source data are provided in Source Data Extended Data Fig. 7.

**Extended Data Figure 8:**
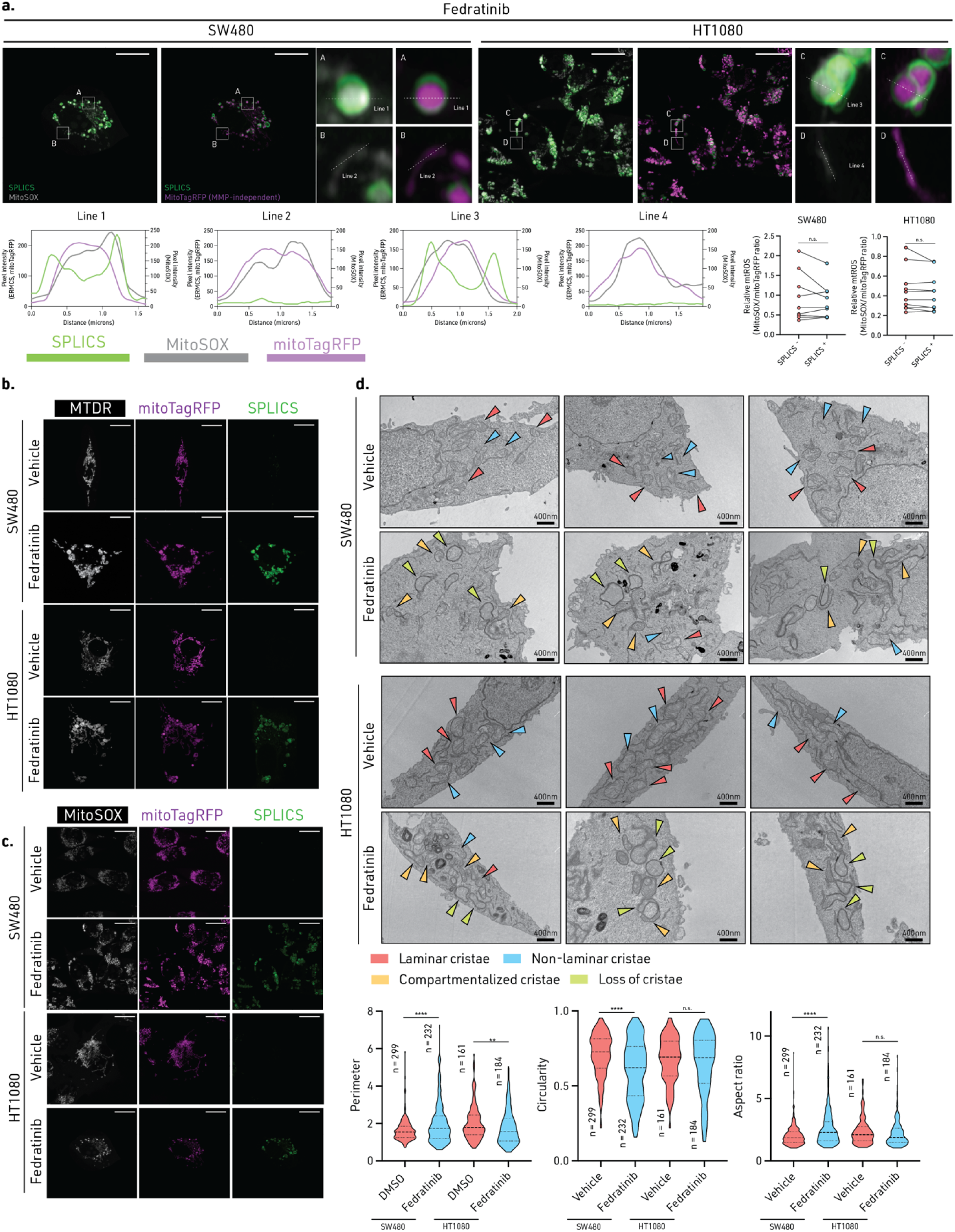
Light and electron microscopy analyses of mitochondria in fedratinib-treated cells. **a**, Confocal images of mtROS (MitoSOX Deep Red) and total mitochondrial mass (mitoTagRFP) from cells treated with vehicle and fedratinib for 24 hr (Upper left). Line intensity profile of representative SPLICS^Hi^ and SPLICS^Lo^ mitochondria (Lower left). Ratiometic analyses of mitochondria membrane potential via MitoSOX/mitoTagRFP ratio were performed on SPLICS^Hi^ and SPLICS^Lo^ mitochondria in cells following treatment with vehicle and fedratinib for 24 hr. Cell number: n = 10 (Right). Scale bar: 1 μm. **b**, Representative confocal images of MitoTracker Deep Red (MTDR) stained cells treated with vehicle and fedratinib for 24 hr. **c**, Representative confocal images of MitoSOX Deep Red stained cells treated with vehicle and fedratinib for 24 hr. **d**, Three representative TEM images in cells treated with vehicle or fedratinib for 24 hr. Cristae structures are annotated in corresponding colors. Red: Laminar cristae. Blue: Non-laminar cristae. Aspect ratio, circularity, and perimeter (μm) of mitochondria in cells treated with vehicle or fedratinib for 24 hr (n=161-299Yellow: Compartmentalized cristae. Green: Loss of cristae. Two-tailed paired t-test were performed on (A). n.s. = not significant. Data are median ± interquartile range (**b**-**d**). Experiments (**a**-**c**) were performed twice, and experiment (**d**) was performed once. n.s. = not significant, **P ≤ 0.01, ***P ≤ 0.001, ****P ≤ 0.0001. Statistical source data are provided in Source Data Extended Data Fig. 8.

**Extended Data Figure 9:**
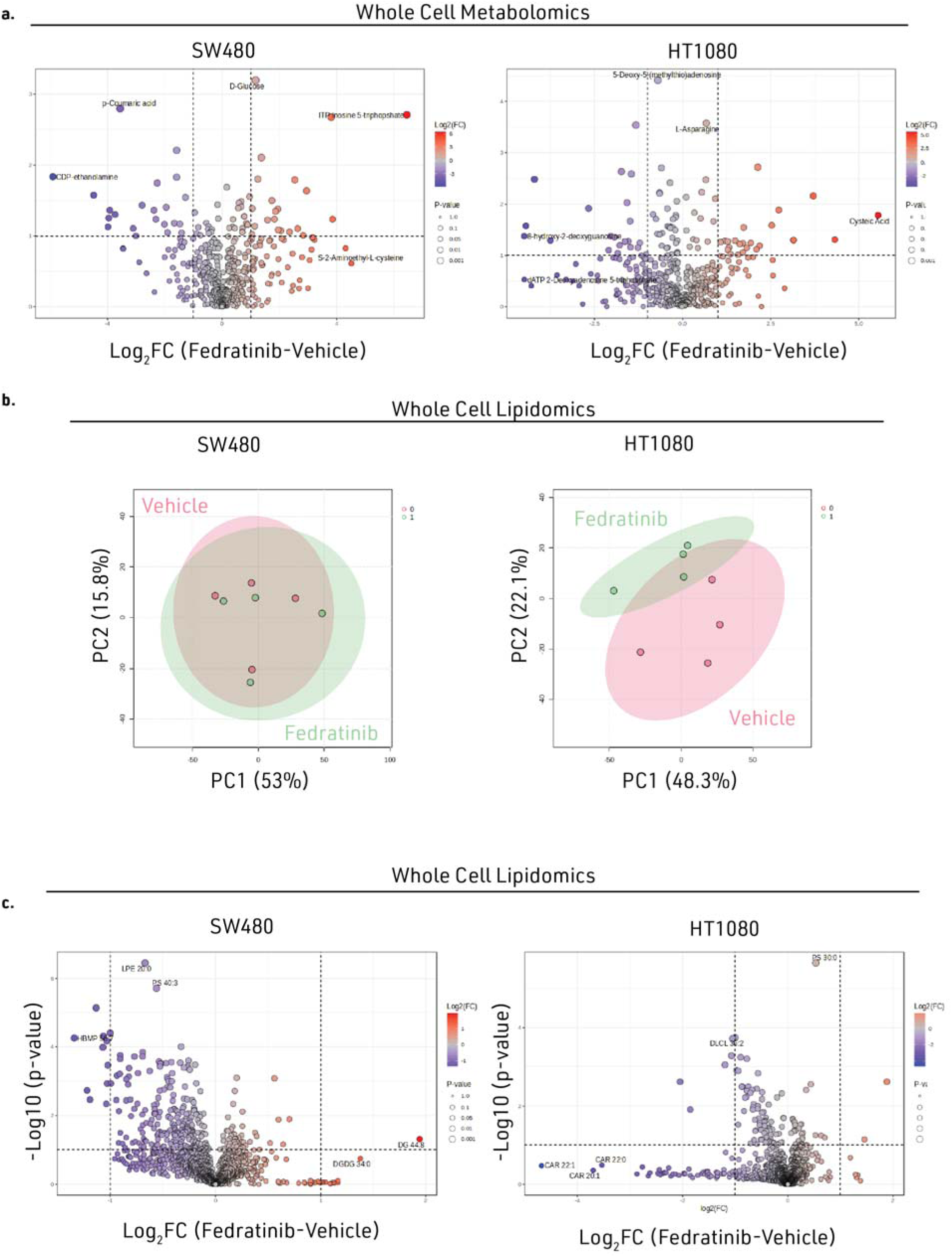
Whole cell metabolomics and lipidomics analyses of fedratinib-treated cells. **a**, Volcano plots of polar metabolites in cells treated with vehicle and fedratinib for 24 hr (Log_2_(Fold change) is plotted with fedratinib-over-vehicle control). **b**, Principal component analysis (PCA) of the lipidome from cells treated with vehicle or fedratinib. **c**, Volcano plots of non-polar lipids from vehicle or fedratinib-treated cells. Log2(FC) is plotted with fedratinib-over-vehicle control. Experiment (**a**) was performed once with three biological replicates. Experiments (**b** and **c**) were performed once with four biological replicates. Metabolomics and Lipidomics raw data is included in Supplementary Table 5 and 6.

**Extended Data Figure 10:**
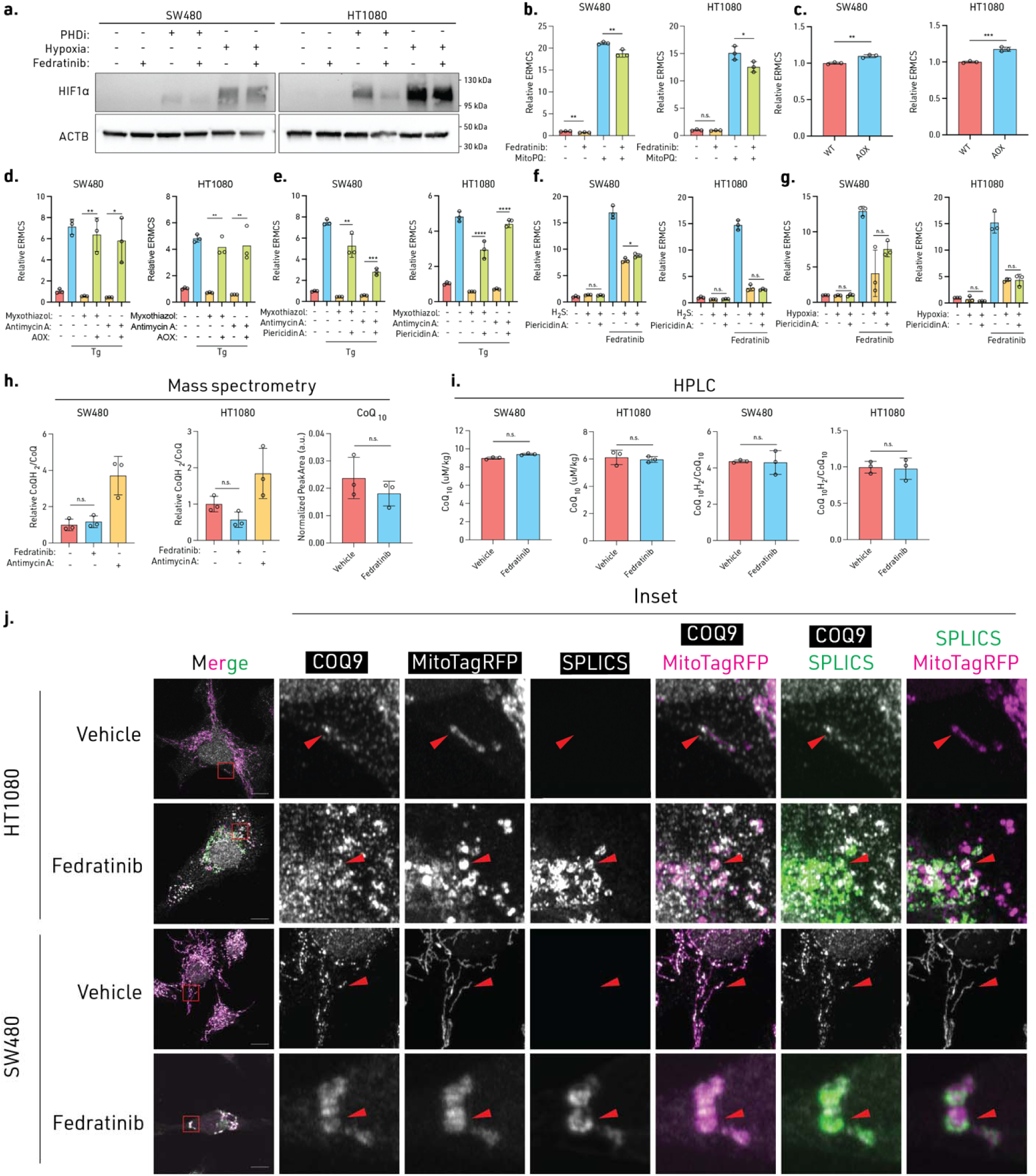
Regulation of ERMCS by mitochondrial ROS inducer and modulators of ETC flow. **a**, Immunoblot of cells treated with PHD inhibitor (FG4592, 50 μM), hypoxia (2% oxygen), and/or fedratinib. **b**, Relative ERMCS in cells treated with mitochondrial-paraquat (mitoPQ, 10 μM) for 24 hr, and then with vehicle or fedratinib for another 24 hr. **c**, Relative ERMCS in AOX expressing cells. **d**, Relative ERMCS in AOX expressing cells treated with complex III inhibitor myxothiazol (50 nM) or antimycin A (100 nM) for 24 hr, and then treated with vehicle or thapsigargin (50 nM) for another 24 hr. **e**, Relative ERMCS in cells treated with complex I and III inhibitor treatment, piericidin A (20 nM), myxothiazol (50 nM), or antimycin A (100 nM) for 24 hr, and then treated with vehicle or thapsigargin (50 nM) for another 24 hr. **f**, Relative ERMCS level in cells co-treated with complex I inhibitor piericidin A (20 nM) and H_2_S or **g**, hypoxia. **h**, Mass spectrometry measurement of relative CoQ redox state from cells treated with vehicle, fedratinib, or positive control antimycin A (100 nM) for 24 hr. Mass spectrometry measurement of CoQ_10_ levels in isolated mitochondria from SW480 cells treated with vehicle or fedratinib for 24 hr. **i**, HPLC measurement of relative CoQ redox state and total CoQ concentrations in cells treated with vehicle or fedratinib for 24 hr. **j**, Confocal imaging of COQ9 in SPLICS/mtRFP cells treated with vehicle or fedratinib for 24 hr. Experiments (**a**-**g**) were performed three times, experiment (**h**, **i**) was performed once with biological triplicates, and experiment (**j**) was performed twice,. Significance was calculated with an unpaired two-tailed t-test. n.s. = not significant, *P ≤ 0.05, **P ≤ 0.01, ***P ≤ 0.005, ****P ≤ 0.001. Data are mean ± SD. Statistical source data are provided in Source Data Extended Data Fig. 9. Immunoblot source data are provided in Supplemental Figure 7.

## See accompanying Microsoft Excel document for Supplementary Tables and movies

**Supplementary Table 1.** Primary results from image-based drug repurposing screen for ERMSC modulators.

**Supplementary Table 2.** Whole cell and FAMS-sorted mitochondrial proteomics results in vehicle (DMSO) treated cells, fedratinib-treated cells. SPLICS^Lo^ and SPLICS^Hi^ mitochondria from vehicle and fedratinib-treated cells.

**Supplementary Table 3.** BrU-seq data for SW480 cells treated with fedratinib for 30 min and 2 hr.

**Supplementary Table 4.** CRISPR screening results for regulators of ERMCS.

**Supplementary Table 5.** Metabolomics information for fedratinib-treated SW480 and HT1080 cells.

**Supplementary Table 6.** Lipidomic information for fedratinib-treated SW480 and HT1080 cells and mitochondria.

**Supplementary Table 7.** Nucleotide sequences used for primers and oligos.

**Supplementary Movie 1:** SW480 cells expressing SPLICS and mitoTagRFP being treated with fedratinib and imaged on Zeiss Lattice Light Sheet 7 for 16 hr.

**Supplementary Movie 2:** HT1080 cells expressing SPLICS and mitoTagRFP being treated with fedratinib and imaged on Zeiss Lattice Light Sheet 7 for 16 hr.

**Supplementary Movie 3:** CLEM Segmentation and EM overlay of SW480 cells treated with DMSO vehicle for 24 hr.

**Supplementary Movie 4:** CLEM Segmentation and EM overlay of SW480 cells treated with fedratinib for 24 hr.

